# Male sex hormone and reduced plakoglobin jointly impair atrial conduction and cardiac sodium currents

**DOI:** 10.1101/2022.06.03.494748

**Authors:** Laura C. Sommerfeld, Andrew P. Holmes, Ting Y. Yu, Christopher O’Shea, Deirdre M. Kavanagh, Jeremy M. Pike, Thomas Wright, Fahima Syeda, Areej Aljehani, Tania Kew, Victor R. Cardoso, S. Nashitha Kabir, Claire Hepburn, Priyanka M. Menon, Sophie Broadway-Stringer, Molly O’Reilly, Anika Witten, Lisa Fortmueller, Susanne Lutz, Alexandra Kulle, Georgios V. Gkoutos, Davor Pavlovic, Wiebke Arlt, Gareth G. Lavery, Richard Steeds, Katja Gehmlich, Monika Stoll, Paulus Kirchhof, Larissa Fabritz

## Abstract

Androgenic anabolic steroids (AAS) are commonly abused by young men. Male sex associates with earlier manifestation of common and rare cardiac conditions including atrial fibrillation and arrhythmogenic right ventricular cardiomyopathy (ARVC). Clinical data suggest an atrial involvement in ARVC. The disease is caused by desmosomal gene defects such as reduced plakoglobin expression. Analysis of clinical records from 146 ARVC patients identified male preponderance and increased prevalence of atrial arrhythmias in patients with definite ARVC. Definite patients displayed ECG changes suggesting atrial remodelling. To study mechanisms of atrial remodelling due to desmosomal vulnerability and AAS, young adult male mice, heterozygously deficient for plakoglobin (Plako^+/-^) and wildtype (WT) littermates, were chronically exposed to 5α-dihydrotestosterone (DHT) or placebo. DHT increased atrial expression of pro-hypertrophic, fibrotic and inflammatory transcripts. DHT caused atrial conduction slowing, decreased peak sodium current density, reduced action potential amplitude and lowered the peak depolarisation rate in Plako^+/-^ but not WT atria. Super-resolution microscopy revealed a reduction in Na_v_1.5 clustering in Plako^+/-^ atrial cardiomyocytes following DHT exposure. These data reveal that AAS combined with plakoglobin deficiency cause pathological atrial electrical remodelling in young male hearts. AAS abuse may increase the risk of atrial myopathy in males with desmosomal gene variants.

## Introduction

Several inherited arrhythmia syndromes develop more severe phenotypes in men carrying pathogenic variants, e.g. Brugada syndrome or Arrhythmogenic Right Ventricular Cardiomyopathy (ARVC) (1). Male sex also associates with greater incidence of atrial arrhythmias, both in the general population and in rare conditions (2–7). Effects mediated by anabolic androgenic steroids (AAS) such as testosterone and the most potent androgen, 5α-dihydrotestosterone (DHT), could contribute. Abuse of AAS is an emerging global health concern, not restricted to elite athletes but common in the general population, with reports indicating a 3.3% lifetime prevalence worldwide (8). AAS are abused predominately by men to increase muscle mass, improve athletic performance and alter appearance (8). However, AAS can cause cardiac pathology, including hypertrophy and electrophysiological changes (9–14). Atrial arrhythmias have recently been associated with elevated total plasma testosterone levels in men (15) and observed in patients known to take AAS devoid of a clinical indication (16–18). Despite these observations, the mechanisms underpinning cardiac electrical remodelling in response to higher levels of AAS are largely unknown.

ARVC has recently been reported to show more adverse outcomes in men (19) related to sex hormone levels (20). Emerging evidence suggests increased incidence of atrial arrhythmias in cardiomyopathies (21). ARVC is often caused by variants in desmosomal genes including plakoglobin (22–25). Plakoglobin is located in desmosomal junctional complexes where it stabilizes cell-cell contacts (26), thereby maintaining mechanical and electrical integrity of the myocardium.

We hypothesized that a substantial proportion of ARVC patients is suffering from clinically relevant atrial arrhythmias and that vulnerability of the desmosome, caused by e.g. plakoglobin reduction, may increase the risk of male sex hormone-induced atrial electrical remodelling.

To test this hypothesis, we screened patient records from definite and non-definite ARVC patients seen at a tertiary center inherited cardiac conditions clinic for atrial arrhythmias. We furthermore employed a murine ARVC model to study the effects of chronically elevated AAS levels in male mice with heterozygous plakoglobin (gamma-catenin) deficiency (Plako^+/-^) (27) and their wildtype (WT) littermates.

## Results

### Atrial arrhythmias and ECG changes in definite ARVC patients

The clinical cohort studied comprised of 146 patients with suspected cardiomyopathy; 97 were identified as “Non-definite” (possible) ARVC cases and 49 as “Definite” ARVC, i.e. presenting with a complete phenotype according to 2010 Task Force Criteria (1). Mean age of the patients at time of ECG analyses was not different between the groups (42 ± 18 years for non-definite vs 43 ± 18 years for definite). 24% of definite ARVC patients experienced atrial fibrillation and/or flutter compared to 3% of the non-definite ARVC patients (Table 1). There was a significant association between sex and ARVC diagnosis type (Table 1, 43% male amongst non-definite vs. 73% male amongst definite patients). Semi-automated analysis of digital ECG lead II recordings focussing on atrial parameters (Figure 1A) showed PR interval prolongation in advanced, definite disease stage (Figure 1B). P wave duration and P wave area were significantly increased in definite compared to non-definite patients. Non-definite ARVC patients exhibited no difference in the analyzed P wave parameters compared to unaffected control subjects. Results of a meta-analysis of clinical observational studies are presented in the discussion.

**Figure 1.**
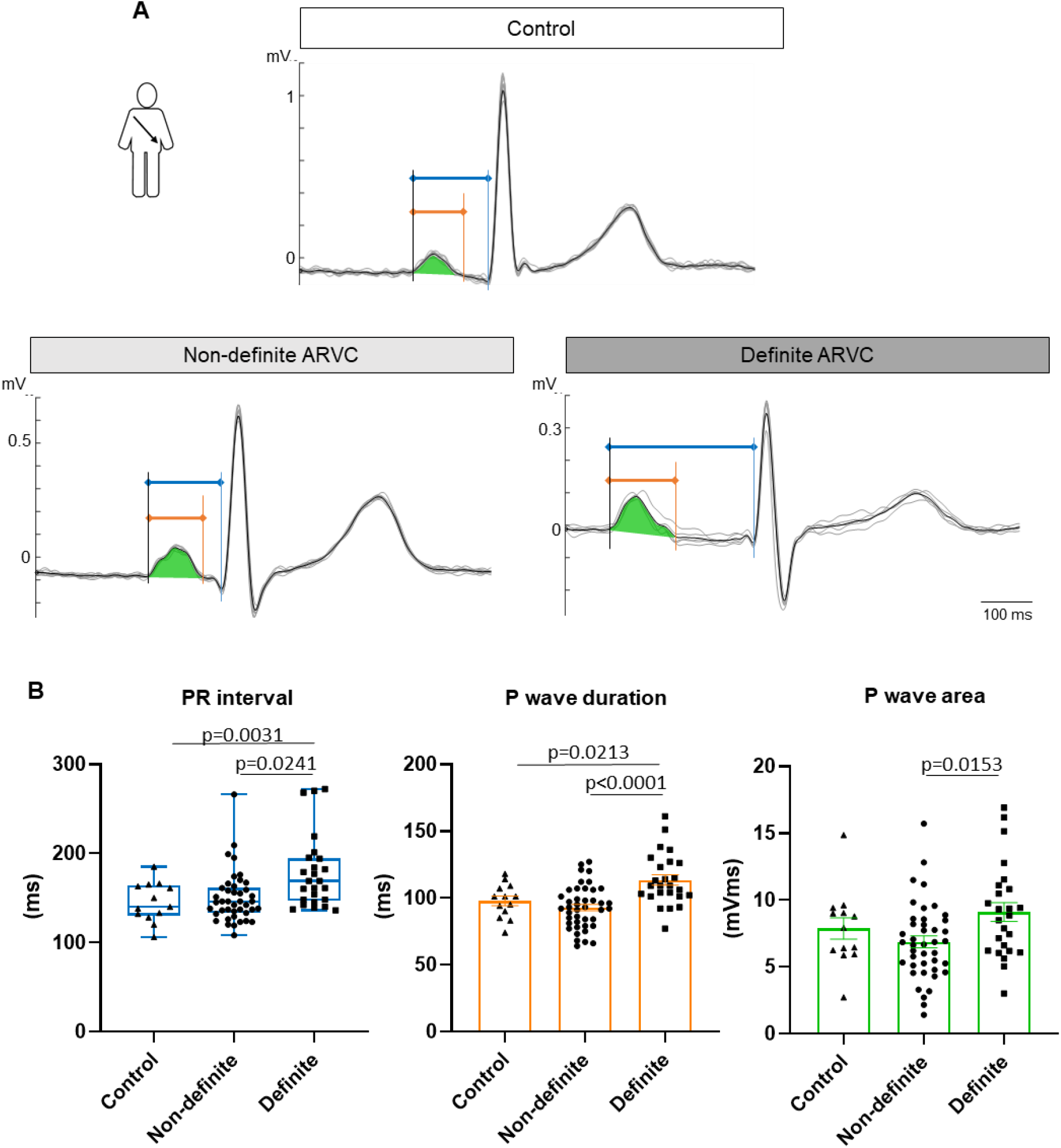
Arrhythmogenic Right Ventricular Cardiomyopathy (ARVC) patients’ and control individuals’ P wave characteristics derived from digital ECG analysis. (**A**) Lead II ECG recordings from an unaffected (control) as well as a non-definite and definite ARVC patient. Individual cardiac cycles over a duration of 10 sec (grey traces) are overlaid by detected R waves and averaged (black trace). PR interval (blue), P wave duration (orange) and P wave area (green) are marked. (**B**) PR interval and P wave characteristics obtained from semi-automated analysis of the averaged ECG. All parameters are derived from lead II recordings. Mean heart rate ± SEM: Control: 73 ± 3 bpm; Non-definite ARVC: 75 ± 2 bpm; Definite ARVC: 60 ± 3 bpm. P-values from post hoc tests are reported on the graphs (Kruskal-Wallis (p<0.05) with Dunn’s post hoc test for PR interval; one-way ANOVA (p<0.05) with Bonferroni post hoc test for P wave duration and area). n (number of patients) = Control: 13, Non-definite ARVC: 42, Definite ARVC: 25

**Table 1.**
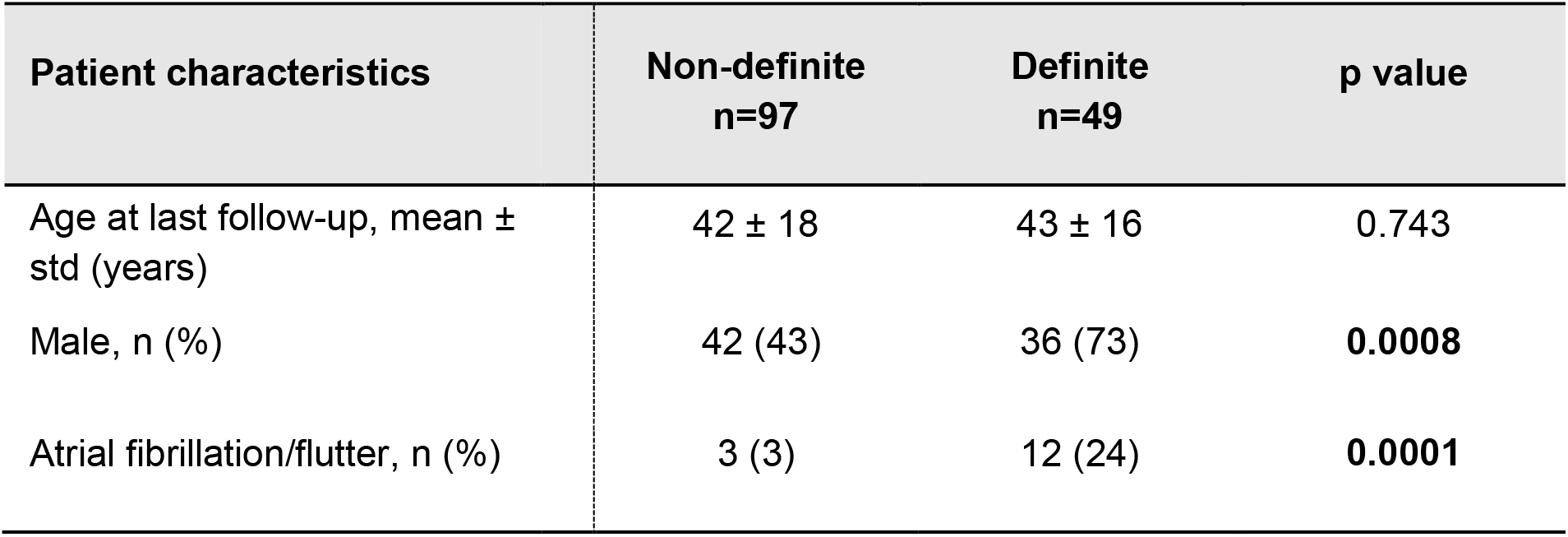
ARVC patient characteristics.

| Patient characteristics | Non-definite<br>n=97 | Definite<br>n=49 | p value |
| --- | --- | --- | --- |
| Age at last follow-up, mean ± std (years) | 42 ± 18 | 43 ± 16 | 0.743 |
| Male, n (%) | 42 (43) | 36 (73) | **0.0008** |
| Atrial fibrillation/flutter, n (%) | 3 (3) | 12 (24) | **0.0001** |

### 5α-dihydrotestosterone (DHT) causes general and cardiac growth response in mice

To study increased androgen exposure and desmosomal instability jointly in atria, Plako^+/-^ mice, an established animal model of ARVC (28) and wildtype (WT) littermates were subjected to chronic DHT treatment over 6 weeks (please refer to Figure 2A for an overview scheme). Treatment led to a 3-4-fold increase in serum DHT concentration in both genotypes compared to controls (Ctr) (Figure 2B). Moreover, it increased body weight, seminal vesicle mass/tibia length ratio and atrial weight/tibia length ratio as well as causing left ventricular hypertrophy in Plako^+/-^ animals (Figure 2C, Supplementary Figure 1).

**Figure 2.**
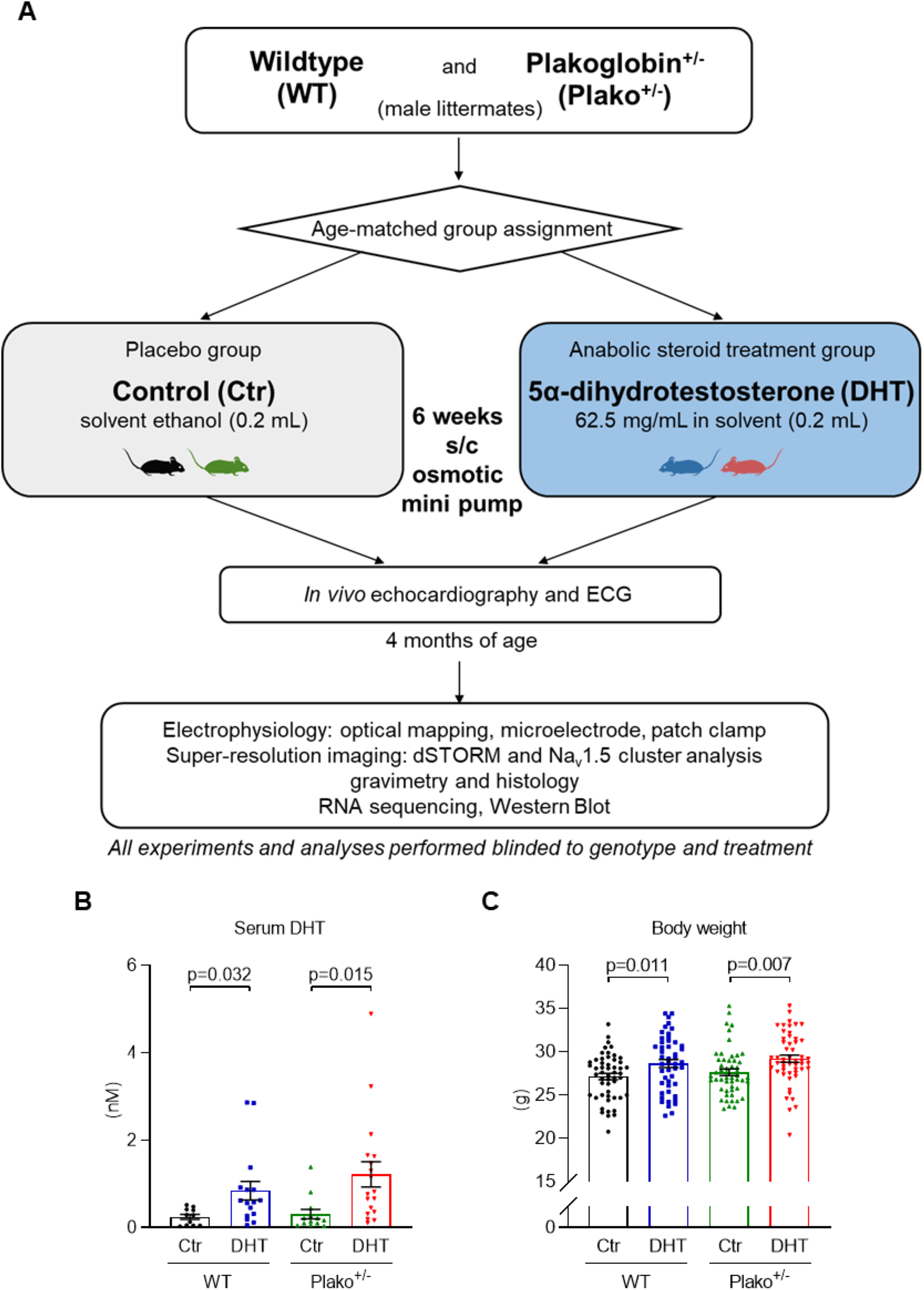
Study design for murine experiments. (**A**) Experimental timeline and methods used in the underlying murine study. All results described were obtained in male WT or Plako^+/-^ mice at 4 months of age, at the end point of the protocol, i.e. after exposure to either 5α-dihydrotestosterone (DHT) or placebo control (Ctr) for 6 weeks. (**B**) Serum DHT concentration (n=11-18 mice/group) and (**C**) body weight (n=48-54 mice/group) are increased following DHT treatment.

### DHT induces ARVC-like atrial ECG changes in plakoglobin-deficient mice

To check whether atrial ECG changes observed in definite ARVC patients are similarly present in the murine model, ECGs were recorded from mice following DHT treatment. Both PR interval as well as P wave duration were prolonged in Plako^+/-^ animals exposed to DHT compared to WT littermates exposed to DHT (Figure 3).

**Figure 3.**
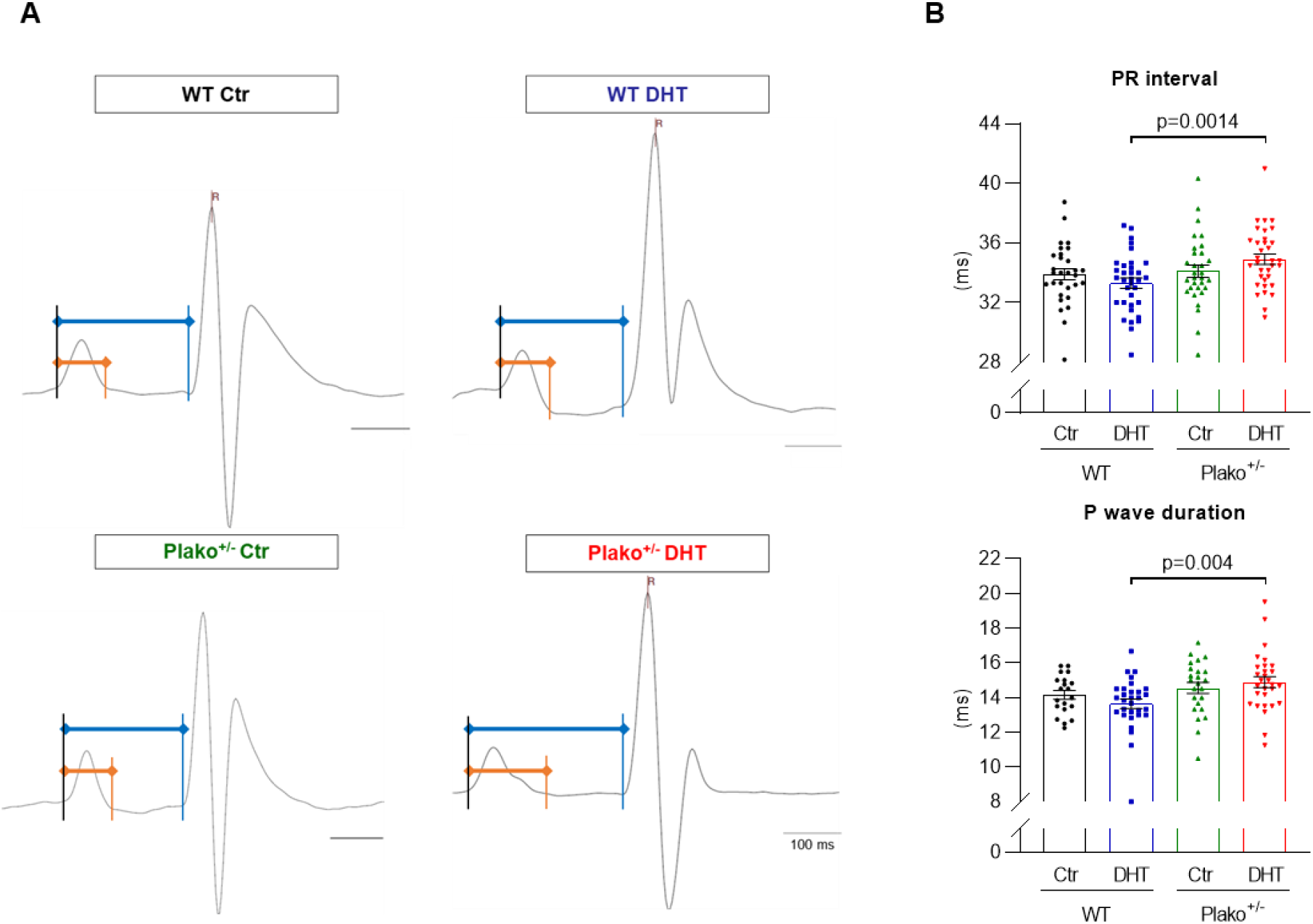
Murine awake ECGs after 6 week chronic control or DHT treatment. (**A**) Exemplary lead II ECG recordings in mice. Shown are compound potentials averaged from 20 subsequent cardiac cycles. PR interval (blue) and monophasic part of P wave (orange) are marked. (**B**) PR interval (n=30-34 mice/group) and P wave duration (n=20-30 mice/group) are prolonged in Plako^+/-^ DHT compared to WT DHT (2-way ANOVA p<0.05 with post-hoc t-test; p-values are indicated on graphs).

### DHT causes atrial unfolded area dilation and atrial conduction slowing in heterozygous plakoglobin-deficient hearts

Left atrial unfolded area to tibia length ratio was increased in Plako^+/-^ DHT compared to Plako^+/-^ Ctr (Figure 4A&B), but not in WT left atria after DHT exposure. Chronic DHT exposure slowed conduction in Plako^+/-^ but not WT atria (Figure 4C&D). The reduction in conduction velocity in Plako^+/-^ DHT left atria was more pronounced at higher pacing frequencies. The Plako^+/-^ left atria exposed to DHT exhibited an overall prolongation of 95% left atrial activation times and increased beat-to-beat activation variability compared to WT Ctr and Plako^+/-^ Ctr (Figure 5). Connexin expression was not impaired (Supplementary Figure 2).

**Figure 4.**
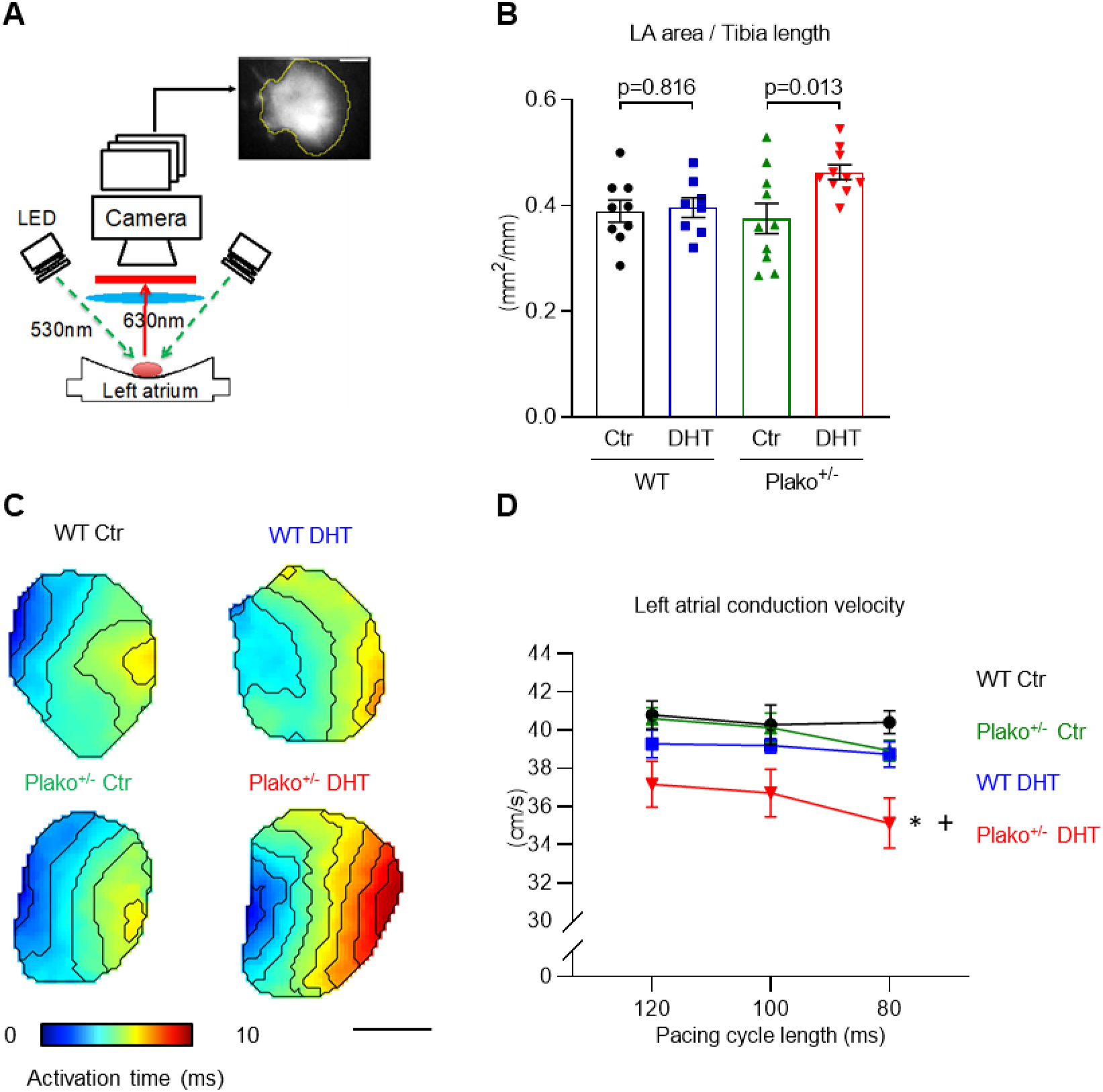
Atrial unfolded area and conduction. (**A**) Experimental setup for optical mapping of isolated left atria (LA), scale bar 1 mm. DHT exposure has a significant effect on (**B**) LA unfolded area (p<0.05, 2-way ANOVA) measured from optical mapping raw images and area is significantly increased in Plako^+/-^ left atria subjected to DHT (post hoc unpaired t-test, p-values indicated on graph, n=8-10 LA per group). (**C**) Exemplary isochronal activation maps of LA at 100 ms pacing cycle length (averaged, scale bar 1 mm). Both, heterozygous deletion of plakoglobin and DHT exposure have a significant effect on (**D**) LA conduction velocity (p<0.05, 2-way repeated measures ANOVA), but it is only significantly decreased in LA of Plako^+/-^ DHT animals (Bonferroni-adjusted post hoc test, +p_adj_<0.05 vs Plako^+/-^ Ctr; *p_adj_ <0.05 vs WT Ctr, across all cycle lengths, n=8-9 LA per group). Data plotted as mean ± SEM.

**Figure 5.**
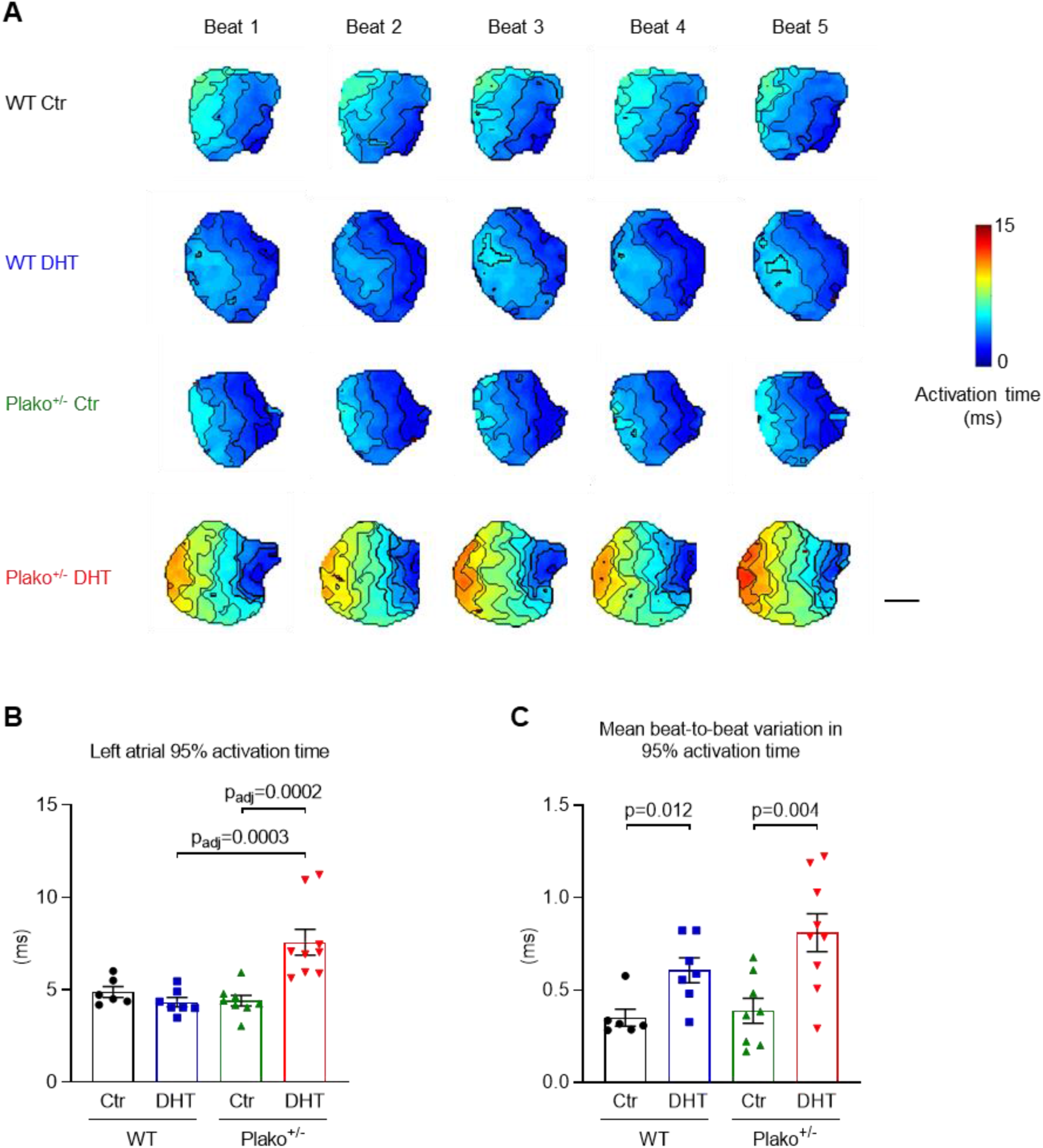
Atrial activation time and beat-to-beat variation. (**A**) Example individual activation maps taken from the final 5 beats of a train of 50 pulses at 80 ms pacing cycle length (CL). DHT exposure has a significant effect on (**B**) beat averaged 95% left atrial activation times. Both genotype and DHT treatment have a significant effect on (**C**) mean beat-to-beat variation in 95% activation times calculated from the final 10 beats of a train of 50 pulses at 80 ms CL (2-way ANOVA with post-hoc analysis as appropriate, p-values indicated). Individual data points denote single left atria (LA). N = WT Control: 6 LA, WT DHT: 7 LA, Plako^+/-^ Control: 8 LA, Plako^+/-^ DHT: 9 LA. Scale bar in (A) indicates 1 mm.

### DHT induces atrial expression of gene profiles implicated in ARVC

Exploratory RNA sequencing analysis confirmed approximately 50% reduction in atrial plakoglobin (*Jup*) expression in Plako^+/-^ compared to WT animals (Figure 6A). In control hearts, gene expression patterns did not markedly differ between genotypes (data not shown). Chronic DHT exposure resulted in significant transcriptional changes in atria of both, WT and Plako^+/-^ mice (Figure 6B&C). DHT activated expression of genes associated with muscle growth (e.g. *Igf1, Mtpn, Myocd*),but additionally also immune (e.g. *C7, Tlr3, Tlr4*) and pro-fibrotic response genes (e.g. *Col1a1, Col3a1, Srf, Lox*) (Figure 6C). Before-mentioned transcriptional changes were similarly induced in left and right atria (Supplementary Figure 3). While atrial cardiac myocyte diameter and endomysial collagen deposition was not significantly different between genotypes after DHT treatment, as quantified in semi-automated histology analysis (Supplementary Figure 5), cell capacitance was increased by DHT treatment in Plako^+/-^ DHT (Figure 8C).

**Figure 6.**
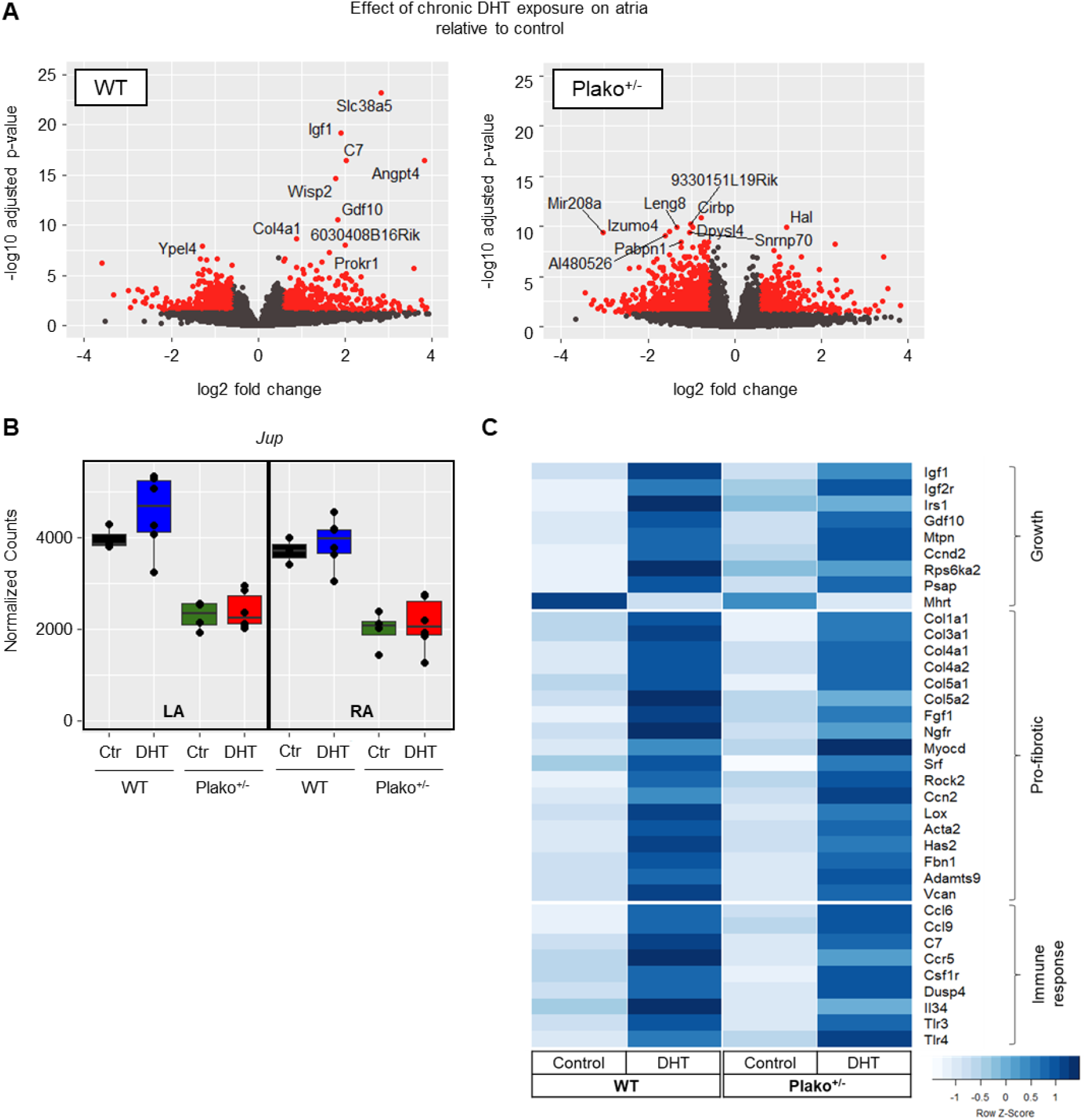
Atrial gene expression profiles. Confirmation of Jup deficiency and growth response. (**A**) Normalized read counts of plakoglobin gene Jup from RNA sequencing analysis from all groups, confirming ~50% reduction of expression in heterozygous knock-out animals (Plako^+/-^) in both left (LA) and right atria (RA) (FDR from DESeq2 WT vs Plako^+/-^ <0.05 for both treatment groups). (**B**) Volcano plot illustrating the effect of DHT treatment, comparing gene expression of WT DHT atria to WT Ctr. Significantly differentially expressed genes at a fold change ≥|0.5| (adjusted p value from DESeq2 < 0.05) are highlighted in red, top 10 hits are labelled. (**C**) Selected genes significantly regulated (adjusted p value from DESeq2 < 0.05) by DHT exposure in at least one of the genotypes. DHT treatment leads to atrial growth, pro-fibrotic and immune response gene expression. n= WT Ctr: 3 LA + 3 RA; WT DHT: 6 LA + 6 RA; Plako^+/-^ Ctr: 4 LA + 4 RA; Plako^+/-^ DHT: 6 LA + 6 RA. Ccnd2, cyclin D2; Gdf10, growth differentiation factor; Igf1, insulin-like growth factor 1; Igf2r, insulin-like growth factor 2 receptor; Irs1, insulin receptor substrate 1; Mhrt, myosin heavy chain associated RNA transcript; Mtpn, myotrophin; Psap, prosaposin; Acta2, smooth muscle (α)-2 actin; Adamts9, ADAM metallopeptidase with thrombospondin type 1 motif 9; Ccn2, connective tissue growth factor; Col1a1/3a1/4a1/4a2/5a1, collagen type 1 alpha 1/3 alpha 1/4 alpha 1/4 alpha 2/5 alpha 1; Fbn1, fibrillin 1; Fgf1, fibroblast growth factor 1; Has2, hyaluronan synthase 2; Lox, lysyl oxidase; Myocd, myocardin; Ngfr, nerve growth factor receptor; Rock2, rho associated coiled-coil containing protein kinase 2; C7, complement C7; Cd, cluster of differentiation 6; Cd9, tetraspanin CD9; Ccr5, C-C motif chemokine receptor 5; Csf1r, colony stimulating factor 1 receptor; Dusp4, dual specificity phosphatase 4; Il34, interleukin 34; Tlr3, toll like receptor 3; Tlr4, toll like receptor 4

**Figure 7.**
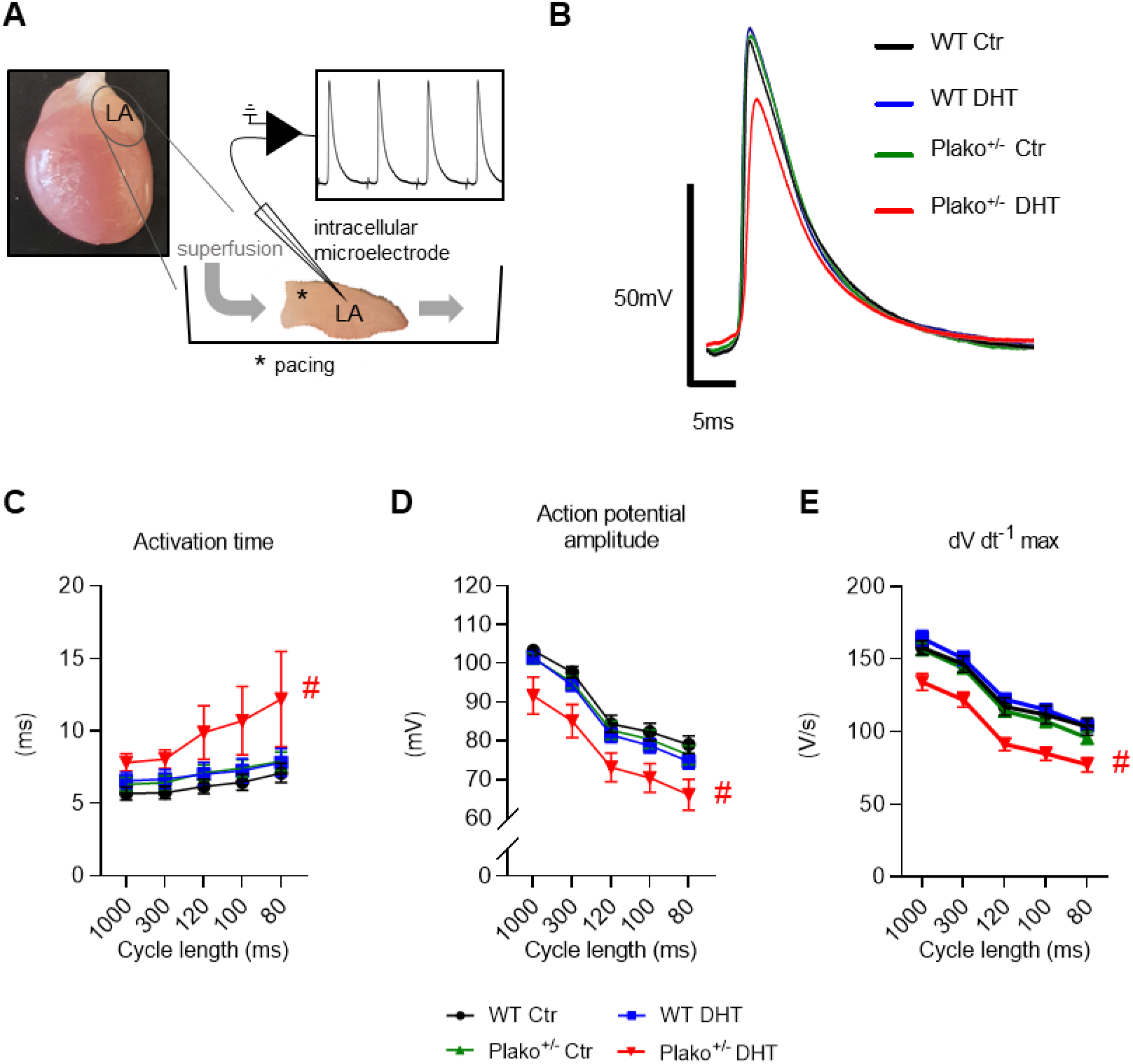
Atrial activation time and action potential characteristics. (**A**) Experimental setup for intracellular microelectrode measurements in paced left atria (LA) to record transmembrane action potentials. Representative examples obtained at 100 ms pacing cycle length are shown in (**B**). Both, heterozygous plakoglobin deletion and DHT exposure have a significant effect on mean (**C**) LA activation time, (**D**) action potential amplitude (APA) and (**E**) maximum upstroke velocity (dV dt^-1^ max) (2-way repeated measures ANOVA, p<0.05). WT Ctr (n=21 cells, N=7 LA), WT DHT (n=13 cells, N=5 LA), Plako^+/-^ Ctr (n=20 cells, N=7 LA) and Plako^+/-^ DHT LA (n=23 cells, N=8 LA). Plako^+/-^ DHT LA have longer activation times, decreased APA and reduced dV dt^-1^ max (Bonferroni-adjusted post hoc analysis: #p_adj_<0.05 vs all other groups, across all cycle lengths). Data averaged per atrium before performing statistical analysis and plotted as mean ± SEM.

**Figure 8.**
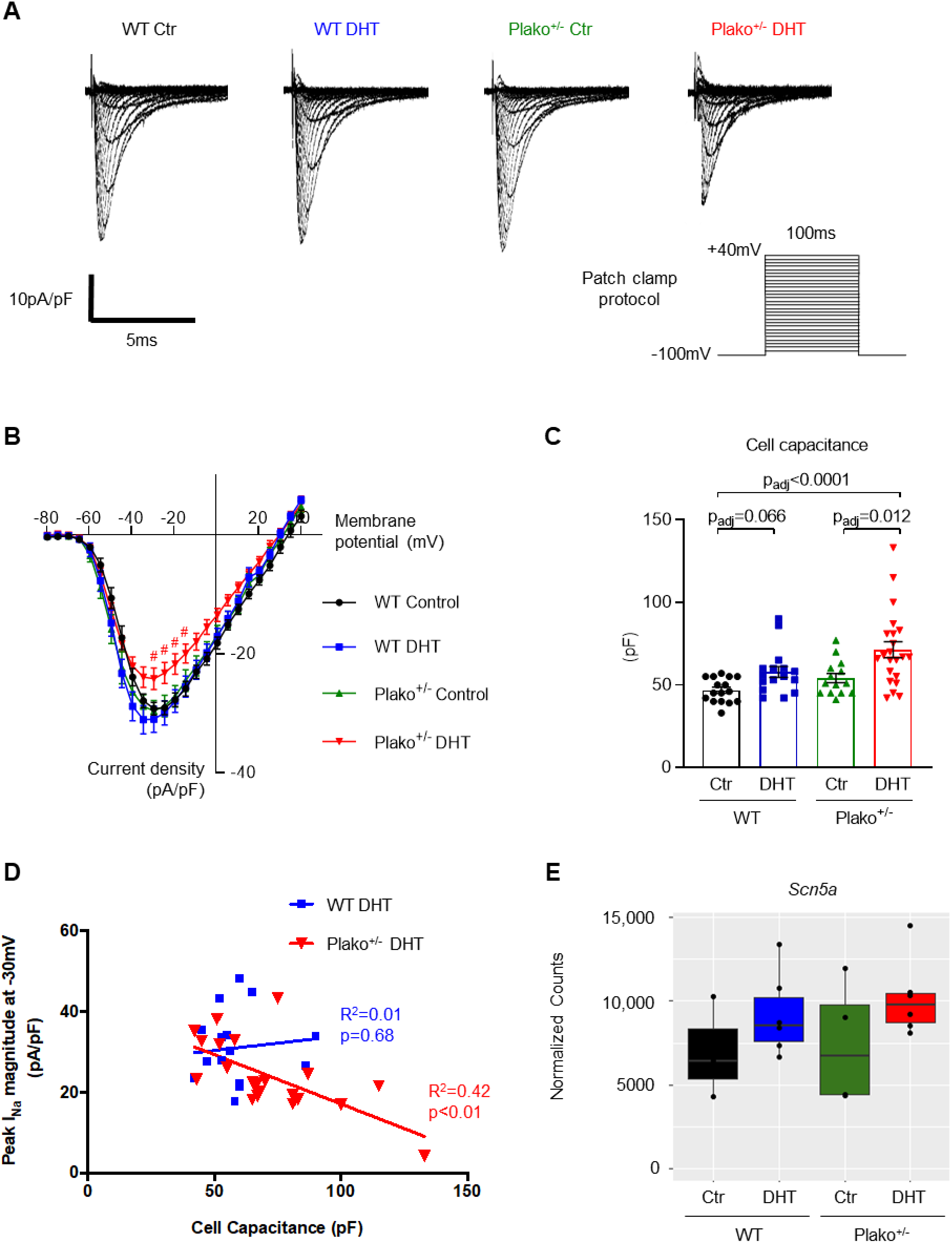
Atrial cardiomyocyte sodium current, cell capacitance and sodium channel expression. (A) Representative left atrial (LA) whole cell peak Na^+^ current (I_Na_) traces measured at test potentials −100 to +40 mV. (**B**) Mean current-voltage relationships for WT Ctr, WT DHT, Plako^+/-^ Ctr and Plako^+/-^ DHT. Plako^+/-^ DHT LA cells have reduced I_Na_ density (2-way repeated measures ANOVA with Bonferroni post hoc analysis: #p_adj_<0.05 vs all other groups at the respective test potential). Data plotted as mean ± SEM. Both genotype and DHT treatment show a significant effect on (**C**) individual cell capacitance (2-way ANOVA, p<0.05) as measured in WT Ctr (n=14 cells, N=5 LA), WT DHT (n=16 cells, N=5 LA), Plako^+/-^ Ctr (n=11 cells, N=4 LA) and Plako^+/-^ DHT (n=21 cells, N=5 LA). Data points shown individually and plotted as mean ± SEM. Bonferroni-corrected post hoc analysis shows a significant increase in cell capacity by DHT compared to Ctr in both genotypes. Cell capacitance is highest in LA cardiomyocytes from Plako^+/-^ DHT. (**D**) Scatter plots of individual LA cell capacitance against peak I_Na_ density for WT Ctr (n=14 cells, N=5 LA), WT DHT (n=16 cells, N=5 LA), Plako^+/-^ Ctr (n=11 cells, N=4 LA) and Plako^+/-^ DHT (n=21 cells, N=5 LA). There is a significant negative correlation of peak I_Na_ density against cell capacitance for Plako^+/-^ but not WT LA cells after DHT treatment; linear regression analysis. (**E**) Normalized read counts of sodium channel transcript *Scn5a* in LA obtained from RNA sequencing analysis (n=3-6 LA per group). Data points shown individually and box whiskers denote IQR, max-min and median.

### DHT reduces action potential amplitude and rate of depolarisation in heterozygous plakoglobin-deficient left atria

The intracellular microelectrode technique was used to record transmembrane action potentials (TAPs) from paced, superfused left atria (Figure 7). Paced left atria isolated from Plako^+/-^ DHT mice showed longer activation times compared to all other groups (Figure 7C, Supplementary Table 1). Action potential amplitude (APA) was reduced in Plako^+/-^ DHT left atria, as well as the peak rate of depolarisation (dV dt^-1^ max) (e.g. 120 ms pacing cycle length WT Ctr: 118±5 V/s; WT DHT: 120±7 V/s; Plako^+/-^ Ctr: 116±4 V/s; Plako^+/-^ DHT: 89±5 V/s) (Figure 7D&E, Supplementary Table 1), both indicative of sodium current impairments. The Plako^+/-^ DHT left atrial cells had a more positive resting membrane potential than WT Ctr (Supplementary Table 1). Chronic DHT exposure did not modify action potential duration (APD) in either genotype (Supplementary Table 1). Beat-averaged left atrial optical APD90, as well as beat-to-beat APD90 variability were similar between all four groups (Supplementary Table 2).

### DHT decreases peak sodium current density in heterozygous plakoglobin-deficient left atrial cardiac myocytes

To examine mechanisms underlying alterations in action potential morphology and conduction in the Plako^+/-^ DHT left atria, whole-cell patch clamp experiments were performed monitoring peak sodium current (I_Na_) amplitude and kinetics. Peak whole-cell I_Na_ density was decreased by approximately 20% in Plako^+/-^ left atrial cells following DHT exposure (Figure 8A&B). Activation kinetics were consistent between all groups (V_50_ activation, WT Ctr: −43±1 mV; WT DHT: −46±1 mV; Plako^+/-^ Ctr: −46±1 mV; Plako^+/-^ DHT: −46±2 mV). DHT exposure tended to cause a left shift in steady-state inactivation kinetics in both WT and Plako^+/-^, being statistically significant in Plako^+/-^ (Supplementary Figure 5A). The 50% recovery time (P50) from inactivation was significantly longer in the Plako^+/-^ DHT, suggestive of a delayed rate of sodium channel recovery (Supplementary Figure 5B&C).

Left atrial cell capacitance was elevated by DHT exposure in Plako^+/-^ (Figure 8C). Of note, there was a considerable spread of individual cell capacitance in DHT-exposed groups, indicative of variable degrees of hypertrophy. I_Na_ density was negatively correlated against cell capacitance in Plako^+/-^ left atria cardiac myocytes following DHT treatment (Figure 8D). This negative correlation was not detected in the WT DHT group. These data suggest that a hypertrophic response in the Plako^+/-^ DHT left atrial cardiac myocytes was not matched by a rise in I_Na_, leading to an overall depletion of I_Na_ density. In contrast, hypertrophy in the WT DHT left atria was matched by an increase in I_Na_, so that I_Na_ density was preserved. Whole left atrial tissue RNA expression of sodium voltage gated channel 5 (*Scn5a*) was neither significantly affected by genotype, or DHT exposure (Figure 8E).

### Na_v_1.5 cluster depletion in Plako^+/-^ left atrial cardiac myocytes following DHT exposure

To define a molecular cause of I_Na_ density depletion in Plako^+/-^ DHT left atrial cardiac myocytes, we examined Na_v_1.5 organisation at super-resolution level using direct Stochastic Optical Reconstruction Microscopy (dSTORM, workflow see Supplementary Figure 6). Due to the high variability in T-tubule density between different atrial cells, we focused on Na_v_1.5 channels located within ca. 200 nm of the contact cell surface membrane by using TIRF/HILO. Exemplary super-resolution images of Na_v_1.5 detections, as well as characteristic cluster maps are shown in Figure 9A.

**Figure 9.**
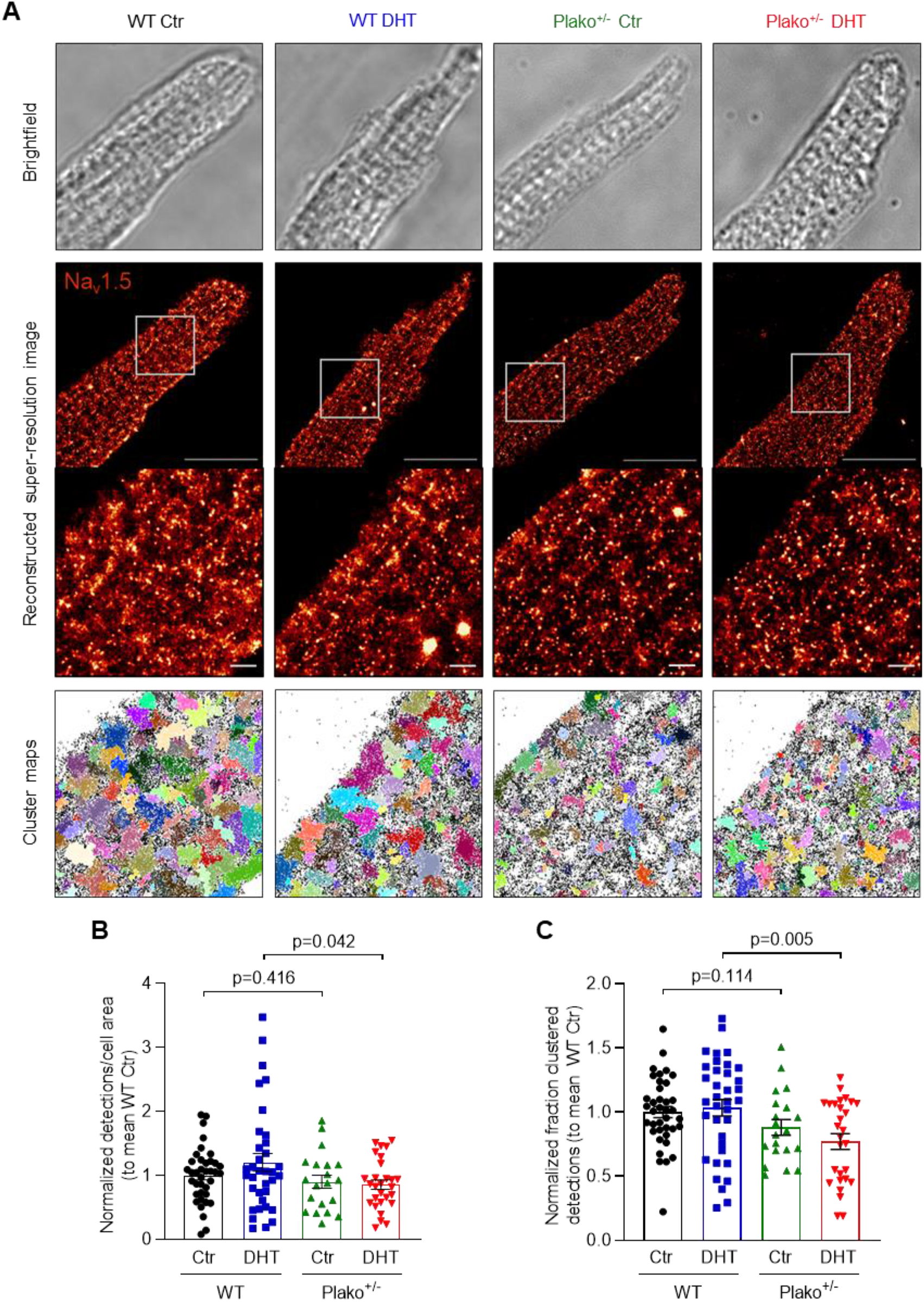
Sodium channel clustering analysis in atrial cardiomyocyte dSTORM images. (**A**) Brightfield images of fixed left atrial (LA) cardiomyocytes prior to dSTORM recording (top lane), reconstructed super-resolution images of membrane Na_v_1.5 and zoom-ins of the indicated area (middle lanes) as well as corresponding cluster maps generated from binary images (bottom lane). Detections allocated to a cluster appear in (arbitrary) colour, non-clustered detections remain black. Scale bar 20 μm and 1.5 μm for zoom-in images, respectively. Genotype has a significant effect on (**B**) number of Na_v_1.5 detections at the LA cardiomyocyte membrane and (**C**) fraction of Na_v_1.5 detections allocated to a cluster (2-way ANOVA, p<0.05). Both number and fraction of clustered membrane Na_v_1.5 is significantly reduced in cardiomyocytes from Plako^+/-^ DHT. Results from post-hoc t-tests are reported on the graphs. For (B) and (C): WT Ctr (n=38 cells, N=7 LA), WT DHT (n=35/36 cells, N=11 LA), Plako^+/-^ Ctr (n=20 cells, N=5 LA) and Plako^+/-^ DHT (n=28 cells, N=6 LA). Data points were normalized to the mean of the respective WT Ctr group on the same imaging session and are shown individually and plotted as mean ± SEM.

Characteristics of identified Na_v_1.5 clusters, i.e. cluster density and cluster area, were similar in all 4 groups (Supplementary Figure 7A&B). The 95% confidence intervals generated from our murine WT Ctr left atria experiments suggest a range of 3-4×10^-3^ Na_v_1.5 channels per nm^2^ cluster (Supplementary Figure 7C, both mean and median at 3×10^-3^ channels per nm^2^ cluster), resulting in an estimated nearest-neighbour distance of 20 nm.

At the Plako^+/-^ DHT left atrial cardiac myocyte membranes, fewer Na_v_1.5 detections were observed compared to WT DHT (Figure 9A&B). Of the total detections, the proportion present in distinct Na_v_1.5 clusters was significantly lower in the Plako^+/-^ DHT compared to WT DHT cardiac myocytes (Figure 9C).

## Discussion

### Main findings

- Atrial arrhythmias and P wave changes are common in patients with ARVC.
- Exposure to the potent androgen 5α-dihydrotestosterone leads to pro-hypertrophic, pro-fibrotic and inflammatory transcriptional signatures in murine atria without overt phenotypic changes.
- Combining chronic 5α-dihydrotestosterone exposure with heterozygous plakoglobin deficiency leads to a profound atrial cardiomyopathy replicating ECG changes in patients with ARVC.
- Mechanistically, increased 5α-dihydrotestosterone concentrations interact with plakoglobin to decrease the number of membrane-localized Na_v_1.5 clusters, reducing atrial sodium current density and causing atrial conduction slowing.

We report that definite ARVC patients exhibit increased P wave area and prolonged PR interval as well as P wave duration on ECG. A phenocopy of these ECG changes is observed in plakoglobin-deficient mice exposed to supraphysiological 5α-dihydrotestosterone (DHT) concentrations. Heterozygous plakoglobin deficiency predisposes cardiac atrial tissue to hypertrophy and a reduction in sodium current density after six weeks of exposure to elevated concentrations of DHT, resulting in a decrease in the peak upstroke velocity of the atrial action potential (dV dt^-1^ max), left atrial conduction slowing and increased electrical beat-to-beat variability. Super-resolution microscopy, dSTORM, identified sarcolemmal depletion of Na_v_1.5 channels and clusters at left atrial cardiac myocytes isolated from Plako^+/-^ DHT-treated mice. This was accompanied by the functional electrophysiological modifications of reduced atrial Na_v_1.5 current density and impaired atrial conduction.

The results underpin a role of plakoglobin in regulating Na_v_1.5 channel cellular localization in the left atrium and in preserving left atrial electrical integrity in response to stressors such as pro-hypertrophic DHT exposure.

### Patients in the definite ARVC disease stage display a high prevalence of atrial arrhythmias, P wave changes on ECG and preponderance of male sex

A quarter (24%) of the patients with definite ARVC had atrial arrhythmias in our cohort. Meta-analysis of published studies (between 1991 and 2021) revealed a weighted mean atrial arrhythmia prevalence of 15% amongst a total of 1915 ARVC patients (Table 3). Over 99% of the reported patients were diagnosed with definite ARVC according to accepted criteria. Male preponderance of atrial phenotypes was reported in other cohorts as well (Table 3). P wave duration and P wave area showed pathological changes in patients with definite ARVC, suggesting pathophysiological remodelling of the atria. P wave prolongation and increased P wave area have been associated with atrial fibrillation (AF) or increased AF recurrence (29–32). Furthermore, definite ARVC patients displayed longer PR intervals compared to non-definite patients and controls, suggesting attenuated conduction of the atrioventricular node. Conduction slowing is a common feature of heart rhythm disorders including AF, and acts by permitting the development and maintenance of both micro and macro re-entry (33). In line with our results, Baturova and colleagues recently found evidence for ARVC disease progression to be paralleled by changes in P wave area (34). Our data substantiate clinical evidence for progressive atrial conduction disturbances with progression of ARVC. Our murine data demonstrate that these changes can arise from an interaction between genetic desmosomal defects and AAS.

**Table 2.**
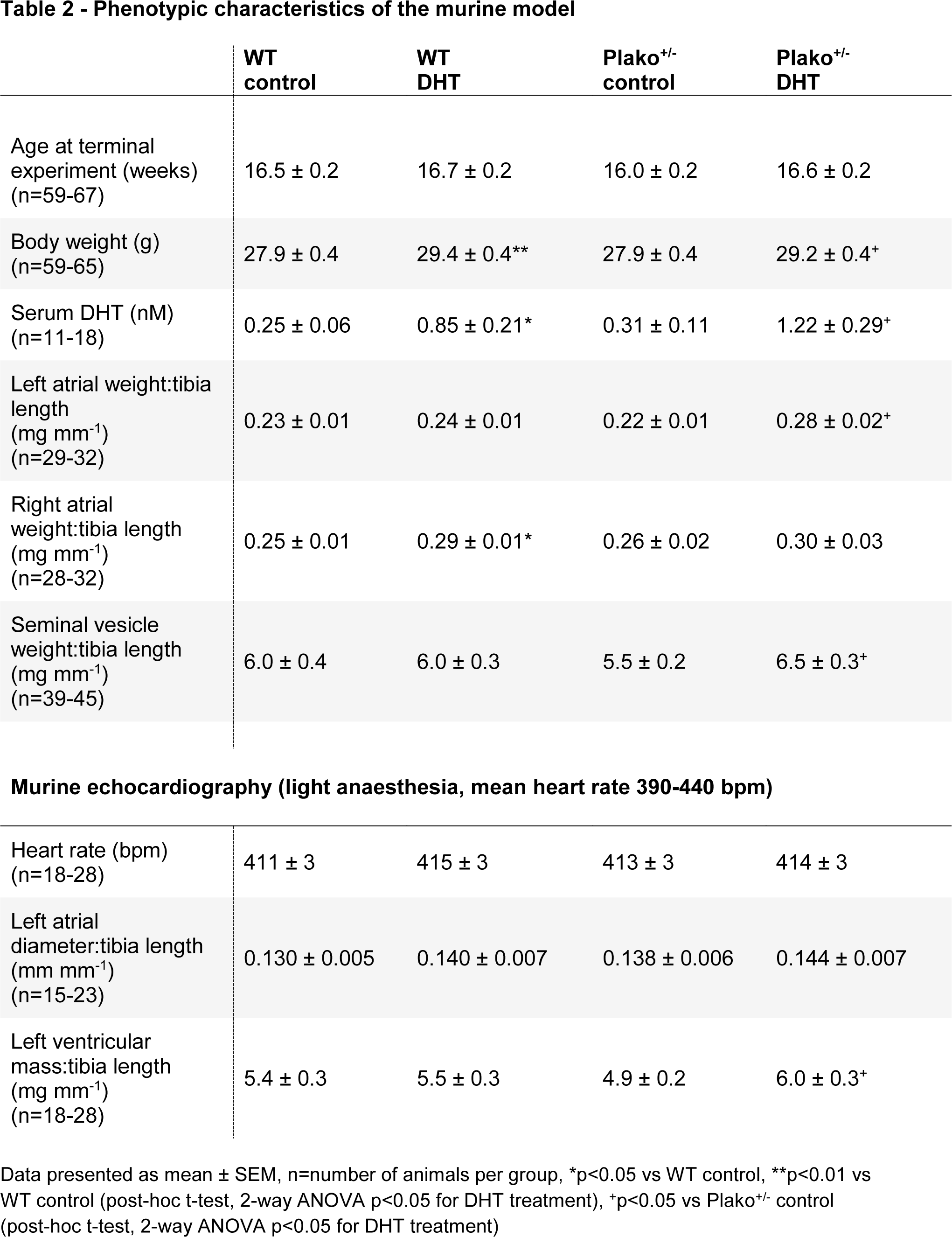
Phenotypic characteristics of the murine model.

|  | WT control | WT DHT | Plako<sup>+/-</sup> control | Plako<sup>+/-</sup> DHT |
| --- | --- | --- | --- | --- |
| Age at terminal experiment (weeks) (n=59-67) | 16.5 ± 0.2 | 16.7 ± 0.2 | 16.0 ± 0.2 | 16.6 ± 0.2 |
| Body weight (g) (n=59-65) | 27.9 ± 0.4 | 29.4 ± 0.4** | 27.9 ± 0.4 | 29.2 ± 0.4<sup>+</sup> |
| Serum DHT (nM) (n=11-18) | 0.25 ± 0.06 | 0.85 ± 0.21* | 0.31 ± 0.11 | 1.22 ± 0.29<sup>+</sup> |
| Left atrial weight:tibia length (mg mm<sup>-1</sup>) (n=29-32) | 0.23 ± 0.01 | 0.24 ± 0.01 | 0.22 ± 0.01 | 0.28 ± 0.02<sup>+</sup> |
| Right atrial weight:tibia length (mg mm<sup>-1</sup>) (n=28-32) | 0.25 ± 0.01 | 0.29 ± 0.01* | 0.26 ± 0.02 | 0.30 ± 0.03 |
| Seminal vesicle weight:tibia length (mg mm<sup>-1</sup>) (n=39-45) | 6.0 ± 0.4 | 6.0 ± 0.3 | 5.5 ± 0.2 | 6.5 ± 0.3<sup>+</sup> |
| **Murine echocardiography (light anaesthesia, mean heart rate 390-440 bpm)** |  |  |  |  |
| Heart rate (bpm) (n=18-28) | 411 ± 3 | 415 ± 3 | 413 ± 3 | 414 ± 3 |
| Left atrial diameter:tibia length (mm mm<sup>-1</sup>) (n=15-23) | 0.130 ± 0.005 | 0.140 ± 0.007 | 0.138 ± 0.006 | 0.144 ± 0.007 |
| Left ventricular mass:tibia length (mg mm<sup>-1</sup>) (n=18-28) | 5.4 ± 0.3 | 5.5 ± 0.3 | 4.9 ± 0.2 | 6.0 ± 0.3<sup>+</sup> |

Data presented as mean ± SEM, n=number of animals per group, \*p<0.05 vs WT control, \*\*p<0.01 vs WT control (post-hoc t-test, 2-way ANOVA p<0.05 for DHT treatment), <sup>+</sup>p<0.05 vs Plako<sup>+/-</sup> control (post-hoc t-test, 2-way ANOVA p<0.05 for DHT treatment)

**Table 3.**
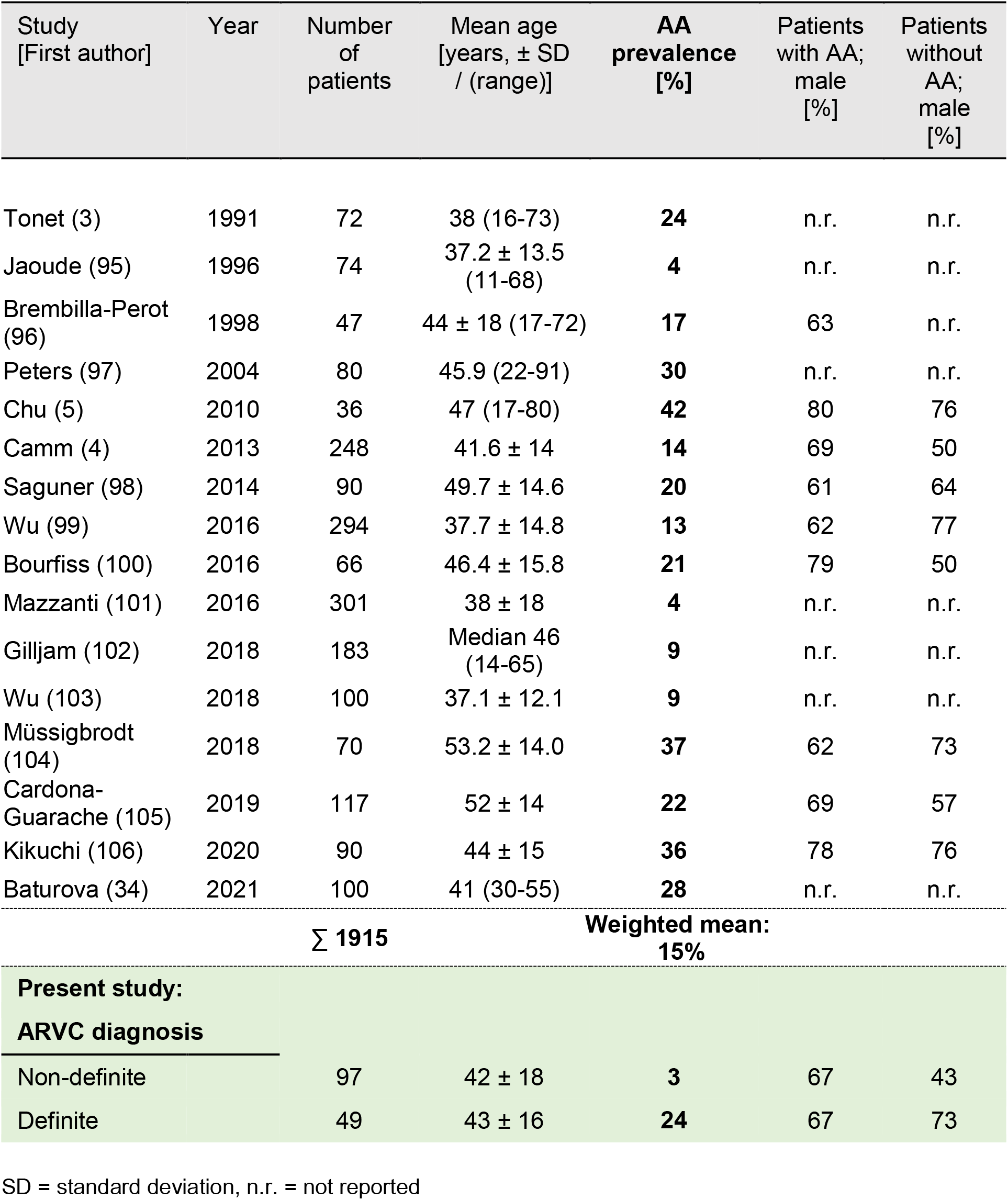
Prevalence of Atrial fibrillation/flutter (Atrial Arrhythmia, AA) in ARVC patients.

### DHT exposure interacts with plakoglobin deficiency leading to atrial electrical dysfunction

Although supraphysiological AAS intake is commonly used by competitive athletes to enhance performance (doping), reports are unsystematic due to underreporting of AAS intake. Murine models represent a unique tool to observe early atrial changes arising in interaction between genetic defects in the desmosome and AAS exposure. We show that chronic DHT exposure combined with heterozygous deletion of plakoglobin prolongs PR interval and P wave duration and slows atrial conduction. Affected atria did not show fibrotic or fatty changes, and connexin expression was not altered. A third important determinant of conduction velocity is the magnitude of depolarising current, primarily carried by Na^+^ through Na_v_1.5 channels (35). Our electrophysiological measurements found reduced atrial action potential amplitude and upstroke velocity (APA and dV dt^-1^ max), demonstrating a decreased sodium current that is sufficient to lower both the magnitude and rate of depolarisation, key contributors to cardiac conduction velocity. A delay of sodium current recovery times, as observed in Plako^+/-^ DHT-treated cardiomyocytes, was enhanced at higher pacing frequencies. Super-resolution microscopy identified reduced availability and altered clustering of sodium channels as a likely mechanism for these functional defects. The reduced availability of Na_v_1.5 channels in cells already operating without a sufficient conduction reserve (36, 37), is likely to cause more pronounced conduction slowing and beat-to-beat variability as observed at the higher pacing frequencies in our experiments.

Supraphysiological DHT concentrations increased atrial expression of genes related to immune response and fibrotic remodelling. Both processes contribute to the pathophysiology of ARVC as well as AF (38–46) and are likely intertwined. Lysyl oxidase (*Lox*), which we here report to be upregulated in atria in response to DHT, mediates cross-linking of collagen I and collagen III fibrils. Both collagen isoforms, amongst others, were also upregulated in DHT-exposed atria in our study, indicating remodelling of the extracellular matrix (ECM). Lox overexpression in mice was also found to accelerate the inflammatory response during angiotensin II (AngII)-induced cardiac hypertrophy, including increased cytokine levels (47). Among the immune-response genes increased in atrial expression upon DHT treatment were the Toll-like receptor 3 (*Tlr3*) and the Toll-like receptor 4 (*Tlr4*) in our study. Silencing or deficiency of these receptors has been demonstrated to improve cardiac function post myocardial infarction in rodent models by attenuating inflammatory cytokine production and fibrotic scar formation (48, 49). Toll-like receptors can be activated by endogenous ligands, including components of the extracellular matrix (50), and have been associated with matrix turn-over.

Excessive ECM deposition is another pathological driver of conduction defects (51, 52) and increased perivascular fibrosis has been observed in the ventricular myocardium of male rats in response to 20 days of testosterone administration (53). Rho associated coiled-coil containing protein kinase 2 (*Rock2*)-dependent pathways, implicated in cardiac fibroblast activation, can regulate expression of ECM component levels, including CTGF (*Ccn2*) and α-smooth muscle actin (*Acta2*) (54). Indeed, the expression of *Rock2, Ccn2*, as well as *Acta2* transcript, was upregulated in atria after chronic DHT exposure. *Ccn2* expression, in response to the pro-hypertrophic stimulus Angll, has previously been shown to be mediated via the transcription factor serum response factor (*Srf*) in cardiac fibroblasts (55), which we also found to be upregulated in atria subjected to high concentrations of DHT. Transcriptional changes did not translate to structural changes other than atrial area dilation, e.g. excessive ECM deposition in atria in our murine study, illustrating the pathophysiologically relevant interaction of DHT-activated profibrotic signalling with desmosomal gene defects for the development of atrial conduction slowing. In the scope of this study, we were able to demonstrate that the combination of a pro-inflammatory/fibrotic environment, caused by DHT, plus the selective reduction in I_Na_ in Plako^+/-^ cardiomyocytes is sufficient to induce conduction slowing and increased beat-to-beat heterogeneity in these vulnerable atria.

While this is the first study showing the molecular effect on *atria*, it has been previously demonstrated that androgens, including DHT, induce a hypertrophic response in *ventricular* cardiomyocytes (12). One of the genes consistently elevated in atrial expression after DHT exposure was insulin growth factor 1 (*Igf1*), a known driver of (cardiac) muscle growth (56, 57). *Igf1* mRNA expression was also found to contribute to atrial fibrotic remodelling and AF inducibility in a rodent model (58). *IGF1*, containing an androgen response element within its promoter region (59), is a well-established target of androgen-receptor mediated gene activation. To our knowledge, we are the first to show that supraphysiological plasma DHT concentration can induce upregulation of *Igf1* transcript levels in atria. Further evidence of the pro-hypertrophic atrial gene response to DHT exposure can be based on the downregulation of the lncRNA Myosin heavy chain associated RNA transcript (*Mhrt*). Repression of *Mhrt* expression has previously been established as a prerequisite for stress-induced, pathological cardiac hypertrophy and restoring *Mhrt* expression levels protected murine hearts from pressure overload-induced hypertrophy (60). Mhrt has been shown to inhibit expression of the transcription factor myocardin (*Myocd*) (61) and increased *Myocd* expression in peripheral blood cells of patients has been associated with increased ventricular mass (62). In accordance with this, down-regulation of *Mhrt* expression upon DHT exposure was paralleled by upregulation of *Myocd* expression compared to control groups. Since we were not able to demonstrate a significant increase in atrial cell diameter in histology at the investigated time point, but were able to demonstrate increased cell capacitance in atrial cardiomyocytes on single cell level as well as atrial area dilation, we propose that the atrial hypertrophic growth response to DHT seen is of an eccentric nature.

### Reduced plakoglobin decreases left atrial Na_v_1.5 cluster availability at the membrane in response to pro-hypertrophic DHT exposure

Super-resolution imaging and electrophysiological techniques suggest that Na_v_1.5 spatial sarcolemmal localization is not random, but rather characterized by formation of distinct clusters (63, 64). To our knowledge, this the first study to evaluate the molecular arrangement of sarcolemma-localized Na_v_1.5 channels in atria through the use of dSTORM to allow for visualisation at nanometer resolution (localisation accuracy estimated to be within 20-30 nm). This confirmed that the mismatch between cell size and I_Na_ density following DHT exposure in Plako^+/-^ left atria was due to reduced membrane Na_v_1.5 channel number and/or defective cluster availability. Interestingly, the values we estimate for next-neighbour distance, calculated from cluster density in WT cells, are slightly lower than those reported employing similar super-resolution methods in ventricular cardiac myocytes (64), suggesting slightly higher cluster densities in atrial cardiomyocytes.

The number of plasma membrane-localized Na_v_1.5 detections was reduced in left atrial cardiomyocytes obtained from Plako^+/-^ DHT and showed a similar trend in Plako^+/-^ control. A link between plakoglobin and Na_v_1.5 channel incorporation/trafficking into the cell membrane at the intercalated disc has been reported in ventricles (65, 66). Interactions of desmosomal and intercalated disc proteins with sodium channel complex have been demonstrated in ventricles previously (67) and loss or mutations of the desmosomal components plakophilin-2 and desmoglein-2 were associated with reduced sodium current in different cell and murine models (67–69). A mismatch in cell size and Na_v_1.5 cluster availability could have implications for other cardiomyopathies and pathological hypertrophy (70, 71). Understanding the role of ion channel cluster properties is in its infancy, but our observations are consistent with other recent studies reporting a reduced I_Na_ in response to changes in single-molecule Na_v_1.5 organisation (72, 73).

### Implications for patients and athletes

Our results show that atrial arrhythmias are an important clinical feature of ARVC and confirm male patients are more likely to show a full ARVC phenotype. We demonstrate a previously unknown interaction between defective desmosomal gene expression and exposure to androgenic anabolic steroids (AAS) in atria. This may partially explain the occurrence of atrial conduction slowing and arrhythmias (74–76) in athletes abusing AAS to enhance their performance. Based on our results, searching for desmosomal gene defects in steroid abusers with atrial arrhythmias seems warranted. Such analyses may add to a better understanding of the cardiac damage observed in some of these patients.

Our data, gained from well-controlled murine experiments, demonstrate that reduced expression of plakoglobin, as commonly observed in cardiac tissue of patients with pathogenic mutations in a variety of desmosomal genes (77), renders atria susceptible to AAS-induced pathology. Prevention of atrial arrhythmias in arrhythmogenic cardiomyopathies is of interest also because they can give rise to inappropriate defibrillator shocks in affected patients (4, 78) and compromise cardiac function.

We here add exposure to DHT to the list of stimuli aggravating pro-arrhythmic phenotypes in carriers of desmosomal mutations and demonstrate that this affects atrial electrical function. Our data also provide an explanation for the stronger phenotypic expression in male gene carriers with desmosomal mutations and the observed worsened clinical outcome in ARVC patients with high physiological testosterone levels (20, 22–25).

## Methods

See supplement for full methods.

### Patient record screening for atrial arrhythmias and semi-automated analysis of digital electrocardiograms (ECGs) from ARVC patients

Adult ARVC patients (>18 years of age) seen at a specialty clinic at a tertiary centre between 2010 and 2021 were classified into two disease severity groups: non-definite and definite cases based on 2010 ARVC Task Force Criteria (TFC) (1).

Clinical records were retrospectively reviewed to obtain information from several modalities including imaging, electrophysiology, histopathology, genetic testing and family history, to cumulatively fulfil a diagnostic classification. Patients exhibiting signs of confirmed disease based on specific combinations of minor or major criteria were classified as the “definite” group. Individuals exhibiting signs on diagnostic investigation congruous with the 2010 TFC as “borderline” or “possible” ARVC were cumulatively considered as the “non-definite” group, as they do not confer a confirmed diagnosis of ARVC. Non-definite cases were included in the analysis to represent individuals in the earlier phases of disease, with a less severe profile of phenotypic expression. Atrial fibrillation and flutter status was extracted from previous electrocardiograms (ECG) and clinical letters.

Digital ECG recordings (10 seconds, sampling frequency 500 Hz) from the most recent follow-up in Inherited Cardiac Conditions Clinic were collated and analyzed using Matlab and BioSigKit (https://doi.org/10.21105/joss.00671). To discern the extent of atrial involvement in definite ARVC patients in comparison with non-definite patients as well as controls, family members of index patients without meeting TFC and/or exclusion of ARVC pathogenic variants attained from targeted gene panel testing, were additionally included (“Control”).

Digital ECG analysis was performed by three independent observers in recordings displaying sinus rhythm applying a custom-designed, semi-automated algorithm. ECGs were digitally filtered between 0.5 and 50 Hz and using a Chebyshev type II filter. The R wave was automatically identified, and all complexes within the recording were averaged to improve signal quality. The isoelectric line was defined from P wave start to end.

### Animal husbandry

Wildtype (WT) and plakoglobin deficient (Plako^+/-^) littermate male mice (28), 129/Sv background, were housed in individually ventilated cages, (2-7 mice/ cage), monitored daily under standard conditions: 12 h light/dark cycle, 22±2 °C and 55±10% humidity. Food and water were available *ad libitum*.

### Chronic DHT exposure in the murine model and experimental timeline

Young adult male mice (8-11 weeks) were assigned to either DHT or placebo/control treatment groups in mixed cages and were fitted with subcutaneous osmotic mini-pumps (Alzet 2006), containing either DHT (62.5 mg/mL in ethanol), or solvent alone (Control, Ctr), for 6 weeks (Figure 2). Age and DHT exposure time were matched for all groups. At least 40 minutes before pump implantation, mice were subcutaneously injected with 0.05 mL of Buprenorphine. Pump implant was performed under anaesthesia with isoflurane inhalation (max. 4%) in O2 with a flow rate of 1-2 L/min. Animal handling staff and investigators were blinded to genotype and treatment. Echocardiography and ECG recording was performed at 6 weeks exposure. Murine hearts were then extracted by thoracotomy under deep terminal anaesthesia (4-5% isoflurane in O2, flow rate 1-2 L/min,), and used for *in organ* and *in vitro* experimental analysis.

### Anabolic androgenic steroid measurements

Murine serum DHT concentrations were determined by ultra-performance liquid chromatography-tandem mass spectrometry (LC-MS/MS) (Waters, Milford, MA, USA) as described before (79).

### Murine awake ECG measurements

Murine ECGs were recorded from awake mice using a tunnel system (ecgTunnel, EMKA Technologies, France) as reported previously (80). Analyses were performed on compound potentials averaged from 20 beats taken at three time-points throughout a 5-min recording with comparable heart rates between groups using the EMKA ECG analysis software. Obtained values were then averaged per animal. For P-wave duration the monophasic part of the P-wave was only taken from ECGs displaying a stable isoelectric line.

### Optical mapping of murine left atria

Activation and action potential duration (APD) maps were generated from isolated left atria loaded with the voltage-sensitive dye Di-4-ANEPPS (17.5 μM; Cambridge Bioscience, CA, USA), paced over a range of 120-80 ms cycle length (CL) as described (81–83). To analyze conduction changes in more detail, beat-to-beat variability in whole tissue activation times was evaluated during rapid physiological pacing. To do this, 10 individual activation maps were compared from the final 10 beats of a train of 50 pulses at 80 ms CL (84).

### Murine echocardiography

Echocardiography was performed with a dedicated small animal system (Vevo 2100; Visualsonics Fujifilm, Toronto, Ont, Canada) under light anaesthesia (0.5-2% isoflurane in O2) at a target heart rate of 390-440 bpm (27, 85–87). The investigators were blinded to intervention and genotype. Images were analyzed by a second blinded observer.

### Transmembrane action potential recordings

Transmembrane action potentials were recorded from isolated, superfused murine left atria as described previously (81–83, 88).

### Cellular electrophysiology

Individual murine left atrial cardiac myocytes were isolated by perfusion with a Tyrode’s enzyme solution containing 20 μg/mL Liberase™ (Roche, Indianapolis, IN) or a collagenase+protease mix, 20 mM taurine and 30 μM CaCl2, via Langendorff over a period of 10-15 min (81, 83). I_Na_ were evoked in voltage-clamp mode using standard protocols and low Na^+^ solution (89). All currents were normalized to cell capacitance.

### Murine atrial fiber size and collagen composition

10 μm left atrial transverse sections were stained with FITC-conjugated wheat germ agglutinin (lectin) and cell diameter as well as endomysial fibrosis were quantified using an automated analysis tool (90).

### Super-resolution microscopy and Na_v_1.5 cluster analysis

Freshly isolated murine left atrial cardiac myocytes were plated on 10mm diameter laminin-coated coverslips (Mattek, 35 mm dish, 1.5# coverglass), fixed, permeabilized and blocked. Cells were stained with primary rabbit anti-Na_v_1.5 antibody (ASC-005, 1:50, Alomone Laboratories, Jerusalem, Israel) and after additional washing and blocking they were incubated in secondary antibody (F(ab’)2-Goat anti-Rabbit IgG, Alexa Fluor 647, A212-56, 1:1000, ThermoFisher Scientific, Waltham, MA, USA). Direct stochastic optical reconstruction microscopy (dSTORM) (91) experiments were performed on a NIKON Eclipse Ti inverted N-STORM microscope equipped with a NIKON APO 100 x 1.49 NA total internal reflection fluorescence (TIRF) oil immersion objective. Immunolabelled samples were imaged in 0.5 mg/mL glucose oxidase, 40 μg/mL catalase, 10% wt/vol glucose and 100 mM MEA in PBS, pH 7.4 to induce Alexa 647 blinking. During dSTORM acquisition, the sample was continuously illuminated at 640 nm for 20,000 frames. Final rendered images of the localized molecules were generated using ThunderSTORM (92) and false coloured in FIJI. Final detections were subjected to a segmentation protocol, using a persistence-based clustering approach (93).

### RNA sequencing of murine left atria

Poly(A) mRNA-enriched libraries were sequenced 75 cycles in a single read mode on the NextSeq-500 System (v2.5 Chemistry, Illumina). Acquired data was trimmed and FASTQ files were aligned using HISAT2 (version 2.1.0; ref Pertea) and the reference genome Ensembl Mus Musculus GRCm38. Differential expression analysis were performed in R (http://www.R-project.org/, R 3.4.1) using the DESeq2 package (94). Differential expression with a false discovery rate <0.05 was deemed significant. Results were visualized using R.

### Statistics

All experiments and analyses were performed blinded to genotype and treatment. Data from murine studies was first subjected to outlier analysis, ROUT method, based around a false discovery rate, where α = 0.01 and outliers were removed (Prism v8, GraphPad Software, La Jolla, CA, USA). Associations between categorical variables were analyzed by Fisher’s exact test. Significance between groups for normally distributed data was taken as p<0.05, ordinary one-way or two-way (repeated) measures ANOVA, with Bonferroni’s post hoc test, as appropriate. Non-normally distributed data was subjected to Kruskal-Wallis with significance level of p<0.05 and Dunn’s post hoc test (Prism v8, GraphPad Software, La Jolla, CA, USA).

### Study approval

Ethical approval for analysis of clinical data from Inherited Cardiac Conditions clinic at the University Hospital Queen Elisabeth, Birmingham, was granted by the local department of research ethics at UHB Hospital (Audit number CARMS-16044).

All animal procedures were approved by the UK Home Office (PPL number 30/2967 and PFDAAF77F) and by the institutional review board of University of Birmingham, UK. All animal procedures conformed to the guidelines from Directive 2010/63/EU of the European Parliament on the protection of animals used for scientific purposes.

## Author contributions

LCS performed murine *in vivo* experiments, dSTORM experiments and analysis, gravimetry and histology analysis, RNAseq analysis, P wave analysis in human ECGs and wrote the manuscript. APH performed and analyzed microelectrode, optical mapping, patchclamp and dSTORM studies, performed gravimetry analysis and wrote the manuscript. TYY and COS set up and performed and analyzed optical mapping experiments. DMK designed dSTORM experiments and trained LCS and APH. JMP generated cluster analysis workflows.TW and PMM performed histology. FS codesigned experiments and trained TW, TYY, SNK. AA and TK screened clinical records according to Task Force Criteria. LCS, TK and COS performed semi-automated patient ECG analysis. CH prepared RNA samples and co-supervised PMM. VRC and GVG analyzed RNAseq. MS and AW performed and analyzed RNAseq. SNK and SBS performed *in vivo* experiments and gravimetry. MOR co-supervised TK. LFo analyzed *in vivo* experiments. SL trained LCS and co-supervised PMM. AK performed mass spectrometry of serum DHT concentrations. WA and GGL advised on AAS experimental design, DP co-supervised COS, RS co-supervised AA, KG co-supervised LCS. PK provided input throughout and co-supervised APH, FS, CH, VRC. LFa (co)-supervised LCS, APH, TYY, COS, TW, FS, AA, TK,VRC, SNK, CH, PMM, SBS, MOR, LFo, designed and coordinated the study and wrote the manuscript. All authors reviewed the results, revised the manuscript and approved the final version of it.

## Acknowledgements

We thank Clara Apicella, Olivia Grech, Pushpa Patel, Genna Riley, and staff of BMSU Birmingham for expert support. We thankfully acknowledge the Centre of Membrane Proteins and Receptors COMPARE (www.birmingham-nottingham.ac.uk/compare) for both their expertise and infrastructure. We thank the Core Facility Genomics of the Medical Faculty Münster, University of Muenster. We thank all members of the Translational Research in Heart Failure and Arrhythmias Cluster for discussion.

This work was partially supported by European Union (grant agreement No 633196 [CATCH ME] to PK and LF), European Union BigData@Heart (grant agreement EU IMI 116074), British Heart Foundation (FS/13/43/30324 to PK and LF; PG/17/30/32961 to PK and APH, PG/20/22/35093 to PK; AA/18/2/34218 to PK and LF; FS/12/40/29712 to KG), German Centre for Cardiovascular Research supported by the German Ministry of Education and Research (DZHK to PK); Leducq Foundation to PK; UOB research development fund (LF). TYY studentship was supported by PSIBS to LF. Funding from the Wellcome Trust was received by KG (201543/B/16/Z) and WA (WT209492/Z/17/Z). KG is also funded by the MRC (MR/V009540/1). The Institute of Cardiovascular Sciences is a recipient of a BHF Accelerator Award (AA/18/2/34218).

GVG acknowledges support from the NIHR Birmingham ECMC, NIHR Birmingham SRMRC, Nanocommons H2020-EU (731032) and the MRC Heath Data Research UK (HDRUK/CFC/01), an initiative funded by UK Research and Innovation, Department of Health and Social Care (England) and the devolved administrations, and leading medical research charities. GVG and WA receive support from the NIHR Birmingham Biomedical Research Centre. The views expressed in this publication are those of the authors and not necessarily those of the NHS, the National Institute for Health and Care Research, the Medical Research Council or the Department of Health.

## References

1. Marcus FI, McKenna WJ, Sherrill D, Basso C, Bauce B, Bluemke DA, et al. Diagnosis of arrhythmogenic right ventricular cardiomyopathy/dysplasia: proposed modification of the Task Force Criteria. European heart journal. 2010;31(7):806–14.

2. Morita H, Kusano-Fukushima K, Nagase S, Fujimoto Y, Hisamatsu K, Fujio H, et al. Atrial fibrillation and atrial vulnerability in patients with Brugada syndrome. J Am Coll Cardiol. 2002;40(8):1437–44.

3. Tonet JL, Castro-Miranda R, Iwa T, Poulain F, Frank R, and Fontaine GH. Frequency of supraventricular tachyarrhythmias in arrhythmogenic right ventricular dysplasia. The American journal of cardiology. 1991;67(13):1153.

4. Camm CF, James CA, Tichnell C, Murray B, Bhonsale A, te Riele AS, et al. Prevalence of atrial arrhythmias in arrhythmogenic right ventricular dysplasia/cardiomyopathy. Heart rhythm. 2013;10(11):1661–8.

5. Chu AF, Zado E, and Marchlinski FE. Atrial arrhythmias in patients with arrhythmogenic right ventricular cardiomyopathy/dysplasia and ventricular tachycardia. The American journal of cardiology. 2010;106(5):720–2.

6. Baturova MA, Haugaa KH, Jensen HK, Svensson A, Gilljam T, Bundgaard H, et al. Atrial fibrillation as a clinical characteristic of arrhythmogenic right ventricular cardiomyopathy: Experience from the Nordic ARVC Registry. Int J Cardiol. 2020;298:39–43.

7. Priori SG, Blomstrom-Lundqvist C, Mazzanti A, Blom N, Borggrefe M, Camm J, et al. 2015 ESC Guidelines for the management of patients with ventricular arrhythmias and the prevention of sudden cardiac death: The Task Force for the Management of Patients with Ventricular Arrhythmias and the Prevention of Sudden Cardiac Death of the European Society of Cardiology (ESC). Endorsed by: Association for European Paediatric and Congenital Cardiology (AEPC). European heart journal. 2015;36(41):2793–867.

8. Sagoe D, Molde H, Andreassen CS, Torsheim T, and Pallesen S. The global epidemiology of anabolic-androgenic steroid use: a meta-analysis and meta-regression analysis. Annals of Epidemiology. 2014;24(5):383–98.

9. Urhausen A, Albers T, and Kindermann W. Are the cardiac effects of anabolic steroid abuse in strength athletes reversible? Heart. 2004;90(5):496–501.

10. Luijkx T, Velthuis BK, Backx FJG, Buckens CFM, Prakken NHJ, Rienks R, et al. Anabolic androgenic steroid use is associated with ventricular dysfunction on cardiac MRI in strength trained athletes. International journal of cardiology. 2013;167(3):664–8.

11. Alizade E, Avci A, Fidan S, Tabakci M, Bulut M, Zehir R, et al. The Effect of Chronic Anabolic-Androgenic Steroid Use on Tp-E Interval, Tp-E/Qt Ratio, and Tp-E/Qtc Ratio in Male Bodybuilders. Ann Noninvasive Electrocardiol. 2015;20(6):592–600.

12. Marsh JD, Lehmann MH, Ritchie RH, Gwathmey JK, Green GE, and Schiebinger RJ. Androgen receptors mediate hypertrophy in cardiac myocytes. Circulation. 1998;98(3):256–61.

13. Medei E, Marocolo M, Rodrigues DD, Arantes PC, Takiya CM, Silva J, et al. Chronic treatment with anabolic steroids induces ventricular repolarization disturbances: Cellular, ionic and molecular mechanism. Journal of molecular and cellular cardiology. 2010;49(2):165–75.

14. Pirompol P, Teekabut V, Weerachatyanukul W, Bupha-Intr T, and Wattanapermpool J. Supra-physiological dose of testosterone induces pathological cardiac hypertrophy. J Endocrinol. 2016;229(1):13–23.

15. Berger D, Folsom AR, Schreiner PJ, Chen LY, Michos ED, O’Neal WT, et al. Plasma total testosterone and risk of incident atrial fibrillation: The Atherosclerosis Risk in Communities (ARIC) study. Maturitas. 2019;125:5–10.

16. Sullivan ML, Martinez CM, and Gallagher EJ. Atrial fibrillation and anabolic steroids. J Emerg Med. 1999;17(5):851–7.

17. Tsai WC, Lee TI, Chen YC, Kao YH, Lu YY, Lin YK, et al. Testosterone replacement increases aged pulmonary vein and left atrium arrhythmogenesis with enhanced adrenergic activity. International journal of cardiology. 2014;176(1):110–8.

18. Lau DH, Stiles MK, John B, Shashidhar, Young GD, and Sanders P. Atrial fibrillation and anabolic steroid abuse. Int J Cardiol. 2007;117(2):e86–7.

19. Rootwelt-Norberg C, Lie OH, Chivulescu M, Castrini AI, Sarvari SI, Lyseggen E, et al. Sex differences in disease progression and arrhythmic risk in patients with arrhythmogenic cardiomyopathy. Europace: European pacing, arrhythmias, and cardiac electrophysiology: journal of the working groups on cardiac pacing, arrhythmias, and cardiac cellular electrophysiology of the European Society of Cardiology. 2021;23(7):1084–91.

20. Akdis D, Saguner AM, Shah K, Wei C, Medeiros-Domingo A, von Eckardstein A, et al. Sex hormones affect outcome in arrhythmogenic right ventricular cardiomyopathy/dysplasia: from a stem cell derived cardiomyocyte-based model to clinical biomarkers of disease outcome. European heart journal. 2017.

21. Wilde AAM, Semsarian C, Marquez MF, Sepehri Shamloo A, Ackerman MJ, Ashley EA, et al. European Heart Rhythm Association (EHRA)/Heart Rhythm Society (HRS)/Asia Pacific Heart Rhythm Society (APHRS)/Latin American Heart Rhythm Society (LAHRS) Expert Consensus Statement on the State of Genetic Testing for Cardiac Diseases. Heart rhythm. 2022.

22. Asimaki A, Syrris P, Wichter T, Matthias P, Saffitz JE, and McKenna WJ. A novel dominant mutation in plakoglobin causes Arrhythmogenic right ventricular cardiomyopathy. American journal of human genetics. 2007;81(5):964–73.

23. Protonotarios N, Tsatsopoulou A, Anastasakis A, Sevdalis E, McKoy G, Stratos K, et al. Genotype-phenotype assessment in autosomal recessive arrhythmogenic right ventricular cardiomyopathy (Naxos disease) caused by a deletion in plakoglobin. Journal of the American College of Cardiology. 2001;38(5):1477–84.

24. Antoniades L, Tsatsopoulou A, Anastasakis A, Syrris P, Asimaki A, Panagiotakos D, et al. Arrhythmogenic right ventricular cardiomyopathy caused by deletions in plakophilin-2 and plakoglobin (Naxos disease) in families from Greece and Cyprus: genotype-phenotype relations, diagnostic features and prognosis. European heart journal. 2006;27(18):2208–16.

25. McKoy G, Protonotarios N, Crosby A, Tsatsopoulou A, Anastasakis A, Coonar A, et al. Identification of a deletion in plakoglobin in arrhythmogenic right ventricular cardiomyopathy with palmoplantar keratoderma and woolly hair (Naxos disease). Lancet (London, England). 2000;355(9221):2119–24.

26. Li J, Swope D, Raess N, Cheng L, Muller EJ, and Radice GL. Cardiac tissue-restricted deletion of plakoglobin results in progressive cardiomyopathy and activation of {beta}-catenin signaling. Mol Cell Biol. 2011;31(6):1134–44.

27. Fabritz L, Hoogendijk MG, Scicluna BP, van Amersfoorth SC, Fortmueller L, Wolf S, et al. Load-reducing therapy prevents development of arrhythmogenic right ventricular cardiomyopathy in plakoglobin-deficient mice. J Am Coll Cardiol. 2011;57(6):740–50.

28. Kirchhof P, Fabritz L, Zwiener M, Witt H, Schafers M, Zellerhoff S, et al. Age- and training-dependent development of arrhythmogenic right ventricular cardiomyopathy in heterozygous plakoglobin-deficient mice. Circulation. 2006;114(17):1799–806.

29. Weinsaft JW, Kochav JD, Kim J, Gurevich S, Volo SC, Afroz A, et al. P wave area for quantitative electrocardiographic assessment of left atrial remodeling. PLoS One. 2014;9(6):e99178.

30. Magnani JW, Zhu L, Lopez F, Pencina MJ, Agarwal SK, Soliman EZ, et al. P-wave indices and atrial fibrillation: cross-cohort assessments from the Framingham Heart Study (FHS) and Atherosclerosis Risk in Communities (ARIC) study. Am Heart J. 2015;169(1):53–61 e1.

31. Soliman EZ, Prineas RJ, Case LD, Zhang ZM, and Goff DC, Jr. Ethnic distribution of ECG predictors of atrial fibrillation and its impact on understanding the ethnic distribution of ischemic stroke in the Atherosclerosis Risk in Communities (ARIC) study. Stroke; a journal of cerebral circulation. 2009;40(4):1204–11.

32. Gorenek B, Birdane A, Kudaiberdieva G, Goktekin O, Cavusoglu Y, Unalir A, et al. P wave amplitude and duration may predict immediate recurrence of atrial fibrillation after internal cardioversion. Ann Noninvasive Electrocardiol. 2003;8(3):215–8.

33. Kirchhof P. The future of atrial fibrillation management: integrated care and stratified therapy. Lancet. 2017;390(10105):1873–87.

34. Baturova MA, Svensson A, Aneq MA, Svendsen JH, Risum N, Sherina V, et al. Evolution of P-wave indices during long-term follow-up as markers of atrial substrate progression in arrhythmogenic right ventricular cardiomyopathy. Europace: European pacing, arrhythmias, and cardiac electrophysiology: journal of the working groups on cardiac pacing, arrhythmias, and cardiac cellular electrophysiology of the European Society of Cardiology. 2021;23(23 Suppl 1):i29–i37.

35. Heijman J, Guichard JB, Dobrev D, and Nattel S. Translational Challenges in Atrial Fibrillation. Circulation research. 2018;122(5):752–73.

36. van Rijen HV, and de Bakker JM. Penetrance of monogenetic cardiac conduction diseases. A matter of conduction reserve? Cardiovasc Res. 2007;76(3):379–80.

37. van Rijen HV, de Bakker JM, and van Veen TA. Hypoxia, electrical uncoupling, and conduction slowing: Role of conduction reserve. Cardiovasc Res. 2005;66(1):9–11.

38. Campian ME, Verberne HJ, Hardziyenka M, de Groot EA, van Moerkerken AF, van Eck-Smit BL, et al. Assessment of inflammation in patients with arrhythmogenic right ventricular cardiomyopathy/dysplasia. Eur J Nucl Med Mol Imaging. 2010;37(11):2079–85.

39. Chelko SP, Asimaki A, Lowenthal J, Bueno-Beti C, Bedja D, Scalco A, et al. Therapeutic Modulation of the Immune Response in Arrhythmogenic Cardiomyopathy. Circulation. 2019.

40. Frustaci A, Chimenti C, Bellocci F, Morgante E, Russo MA, and Maseri A. Histological substrate of atrial biopsies in patients with lone atrial fibrillation. Circulation. 1997;96(4):1180–4.

41. Chung MK, Martin DO, Sprecher D, Wazni O, Kanderian A, Carnes CA, et al. C-reactive protein elevation in patients with atrial arrhythmias: inflammatory mechanisms and persistence of atrial fibrillation. Circulation. 2001;104(24):2886–91.

42. Aviles RJ, Martin DO, Apperson-Hansen C, Houghtaling PL, Rautaharju P, Kronmal RA, et al. Inflammation as a risk factor for atrial fibrillation. Circulation. 2003;108(24):3006–10.

43. Xu J, Cui G, Esmailian F, Plunkett M, Marelli D, Ardehali A, et al. Atrial extracellular matrix remodeling and the maintenance of atrial fibrillation. Circulation. 2004;109(3):363–8.

44. Boldt A, Wetzel U, Lauschke J, Weigl J, Gummert J, Hindricks G, et al. Fibrosis in left atrial tissue of patients with atrial fibrillation with and without underlying mitral valve disease. Heart. 2004;90(4):400–5.

45. Marcus FI, Fontaine GH, Guiraudon G, Frank R, Laurenceau JL, Malergue C, et al. Right ventricular dysplasia: a report of 24 adult cases. Circulation. 1982;65(2):384–98.

46. Thiene G, Corrado D, Nava A, Rossi L, Poletti A, Boffa GM, et al. Right ventricular cardiomyopathy: is there evidence of an inflammatory aetiology? European heart journal. 1991;12 Suppl D:22–5.

47. Galan M, Varona S, Guadall A, Orriols M, Navas M, Aguilo S, et al. Lysyl oxidase overexpression accelerates cardiac remodeling and aggravates angiotensin II-induced hypertrophy. FASEB J. 2017;31(9):3787–99.

48. Lu C, Ren D, Wang X, Ha T, Liu L, Lee EJ, et al. Toll-like receptor 3 plays a role in myocardial infarction and ischemia/reperfusion injury. Biochimica et biophysica acta. 2014;1842(1):22–31.

49. Liu L, Wang Y, Cao ZY, Wang MM, Liu XM, Gao T, et al. Up-regulated TLR4 in cardiomyocytes exacerbates heart failure after long-term myocardial infarction. J Cell Mol Med. 2015;19(12):2728–40.

50. Okamura Y, Watari M, Jerud ES, Young DW, Ishizaka ST, Rose J, et al. The extra domain A of fibronectin activates Toll-like receptor 4. J Biol Chem. 2001;276(13):10229–33.

51. Verheule S, Sato T, Everett T, Engle SK, Otten D, von der Lohe MR, et al. Increased vulnerability to atrial fibrillation in transgenic mice with selective atrial fibrosis caused by overexpression of TGF-beta 1. Circulation research. 2004;94(11):1458–65.

52. Fabritz L, and Kirchhof P. Selective atrial profibrotic signalling in mice and man. Cardiovasc Res. 2013;99(4):592–4.

53. Papamitsou T, Barlagiannis D, Papaliagkas V, Kotanidou E, and Dermentzopoulou-Theodoridou M. Testosterone-induced hypertrophy, fibrosis and apoptosis of cardiac cells--an ultrastructural and immunohistochemical study. Med Sci Monit. 2011;17(9):BR266–73.

54. Akhmetshina A, Dees C, Pileckyte M, Szucs G, Spriewald BM, Zwerina J, et al. Rho-associated kinases are crucial for myofibroblast differentiation and production of extracellular matrix in scleroderma fibroblasts. Arthritis Rheum. 2008;58(8):2553–64.

55. Ongherth A, Pasch S, Wuertz CM, Nowak K, Kittana N, Weis CA, et al. p63RhoGEF regulates auto-and paracrine signaling in cardiac fibroblasts. J Mol Cell Cardiol. 2015;88:39–54.

56. Kim J, Wende AR, Sena S, Theobald HA, Soto J, Sloan C, et al. Insulin-like growth factor I receptor signaling is required for exercise-induced cardiac hypertrophy. Mol Endocrinol. 2008;22(11):2531–43.

57. Weeks KL, Bernardo BC, Ooi JYY, Patterson NL, and McMullen JR. The IGF1-PI3K-Akt Signaling Pathway in Mediating Exercise-Induced Cardiac Hypertrophy and Protection. Adv Exp Med Biol. 2017;1000:187–210.

58. Wang J, Li Z, Du J, Li J, Zhang Y, Liu J, et al. The expression profile analysis of atrial mRNA in rats with atrial fibrillation: the role of IGF1 in atrial fibrosis. BMC Cardiovasc Disord. 2019;19(1):40.

59. Wu Y, Zhao W, Zhao J, Pan J, Wu Q, Zhang Y, et al. Identification of androgen response elements in the insulin-like growth factor I upstream promoter. Endocrinology. 2007;148(6):2984–93.

60. Han P, Li W, Lin CH, Yang J, Shang C, Nuernberg ST, et al. A long noncoding RNA protects the heart from pathological hypertrophy. Nature. 2014;514(7520):102–6.

61. Luo Y, Xu Y, Liang C, Xing W, and Zhang T. The mechanism of myocardial hypertrophy regulated by the interaction between mhrt and myocardin. Cell Signal. 2018;43:11–20.

62. Kontaraki JE, Parthenakis FI, Patrianakos AP, Karalis IK, and Vardas PE. Altered expression of early cardiac marker genes in circulating cells of patients with hypertrophic cardiomyopathy. Cardiovasc Pathol. 2007;16(6):329–35.

63. Bhargava A, Lin XM, Novak P, Mehta K, Korchev Y, Delmar M, et al. Super-resolution Scanning Patch Clamp Reveals Clustering of Functional Ion Channels in Adult Ventricular Myocyte. Circulation research. 2013;112(8):1112–+.

64. Leo-Macias A, Agullo-Pascual E, Sanchez-Alonso JL, Keegan S, Lin XM, Arcos T, et al. Nanoscale visualization of functional adhesion/excitability nodes at the intercalated disc. Nature communications. 2016;7:11.

65. Asimaki A, Kapoor S, Plovie E, Arndt AK, Adams E, Liu ZZ, et al. Identification of a New Modulator of the Intercalated Disc in a Zebrafish Model of Arrhythmogenic Cardiomyopathy. Sci Transl Med. 2014;6(240):15.

66. Noorman M, Hakim S, Kessler E, Groeneweg JA, Cox M, Asimaki A, et al. Remodeling of the cardiac sodium channel, connexin43, and plakoglobin at the intercalated disk in patients with arrhythmogenic cardiomyopathy. Heart rhythm: the official journal of the Heart Rhythm Society. 2013;10(3):412–9.

67. Rizzo S, Lodder EM, Verkerk AO, Wolswinkel R, Beekman L, Pilichou K, et al. Intercalated disc abnormalities, reduced Na(+) current density, and conduction slowing in desmoglein-2 mutant mice prior to cardiomyopathic changes. Cardiovascular research. 2012;95(4):409–18.

68. Sato PY, Musa H, Coombs W, Guerrero-Serna G, Patino GA, Taffet SM, et al. Loss of plakophilin-2 expression leads to decreased sodium current and slower conduction velocity in cultured cardiac myocytes. Circ Res. 2009;105(6):523–6.

69. Cerrone M, Lin X, Zhang M, Agullo-Pascual E, Pfenniger A, Chkourko Gusky H, et al. Missense mutations in plakophilin-2 cause sodium current deficit and associate with a Brugada syndrome phenotype. Circulation. 2014;129(10):1092–103.

70. Hofmann F, Fabritz L, Stieber J, Schmitt J, Kirchhof P, Ludwig A, et al. Ventricular HCN channels decrease the repolarization reserve in the hypertrophic heart. Cardiovasc Res. 2012;95(3):317–26.

71. Wagner S, Dybkova N, Rasenack EC, Jacobshagen C, Fabritz L, Kirchhof P, et al. Ca2+/calmodulin-dependent protein kinase II regulates cardiac Na+ channels. J Clin Invest. 2006;116(12):3127–38.

72. Agullo-Pascual E, Lin XM, Leo-Macias A, Zhang ML, Liang FX, Li Z, et al. Super resolution imaging reveals that loss of the C-terminus of connexin43 limits microtubule plus-end capture and Na(V)1.5 localization at the intercalated disc. Cardiovascular research. 2014;104(2):371–81.

73. te Riele A, Agullo-Pascual E, James CA, Leo-Macias A, Cerrone M, Zhang ML, et al. Multilevel analyses of SCN5A mutations in arrhythmogenic right ventricular dysplasia/cardiomyopathy suggest non-canonical mechanisms for disease pathogenesis. Cardiovascular research. 2017;113(1):102–11.

74. Furlanello F, Serdoz LV, Cappato R, and De Ambroggi L. Illicit drugs and cardiac arrhythmias in athletes. Eur J Cardiovasc Prev Rehabil. 2007;14(4):487–94.

75. Akcakoyun M, Alizade E, Gundogdu R, Bulut M, Tabakci MM, Acar G, et al. Long-term anabolic androgenic steroid use is associated with increased atrial electromechanical delay in male bodybuilders. Biomed Res Int. 2014;2014:451520.

76. Nieschlag E, and Vorona E. Doping with anabolic androgenic steroids (AAS): Adverse effects on non-reproductive organs and functions. Rev Endocr Metab Disord. 2015;16(3):199–211.

77. Asimaki A, Tandri H, Huang H, Halushka MK, Gautam S, Basso C, et al. A new diagnostic test for arrhythmogenic right ventricular cardiomyopathy. N Engl J Med. 2009;360(11):1075–84.

78. Takehara N, Makita N, Kawabe J, Sato N, Kawamura Y, Kitabatake A, et al. A cardiac sodium channel mutation identified in Brugada syndrome associated with atrial standstill. J Intern Med. 2004;255(1):137–42.

79. Kulle AE, Riepe FG, Melchior D, Hiort O, and Holterhus PM. A novel ultrapressure liquid chromatography tandem mass spectrometry method for the simultaneous determination of androstenedione, testosterone, and dihydrotestosterone in pediatric blood samples: age-and sex-specific reference data. J Clin Endocrinol Metab. 2010;95(5):2399–409.

80. Silbernagel N, Walecki M, Schafer MK, Kessler M, Zobeiri M, Rinne S, et al. The VAMP-associated protein VAPB is required for cardiac and neuronal pacemaker channel function. FASEB J. 2018;32(11):6159–73.

81. Holmes AP, Yu TY, Tull S, Syeda F, Kuhlmann SM, O’Brien S-M, et al. A Regional Reduction in Ito and IKACh in the Murine Posterior Left Atrial Myocardium Is Associated with Action Potential Prolongation and Increased Ectopic Activity. PloS one. 2016;11(5):e0154077.

82. Yu TY, Syeda F, Holmes AP, Osborne B, Dehghani H, Brain KL, et al. An automated system using spatial oversampling for optical mapping in murine atria. Development and validation with monophasic and transmembrane action potentials. Progress in biophysics and molecular biology. 2014.

83. Syeda F, Holmes AP, Yu TY, Tull S, Kuhlmann SM, Pavlovic D, et al. PITX2 Modulates Atrial Membrane Potential and the Antiarrhythmic Effects of Sodium-Channel Blockers. J Am Coll Cardiol. 2016;68(17):1881–94.

84. O’Shea C, Holmes AP, Yu TY, Winter J, Wells SP, Correia J, et al. ElectroMap: High-throughput open-source software for analysis and mapping of cardiac electrophysiology. Scientific reports. 2019;9(1):1389.

85. Kirchhof P, Kahr PC, Kaese S, Piccini I, Vokshi I, Scheld HH, et al. PITX2c is expressed in the adult left atrium, and reducing Pitx2c expression promotes atrial fibrillation inducibility and complex changes in gene expression. Circ Cardiovasc Genet. 2011;4(2):123–33.

86. Blana A, Kaese S, Fortmuller L, Laakmann S, Damke D, van Bragt K, et al. Knock-in gain-of-function sodium channel mutation prolongs atrial action potentials and alters atrial vulnerability. Heart Rhythm. 2010;7(12):1862–9.

87. Fabritz L, Kirchhof P, Fortmuller L, Auchampach JA, Baba HA, Breithardt G, et al. Gene dose-dependent atrial arrhythmias, heart block, and brady-cardiomyopathy in mice overexpressing A(3) adenosine receptors. Cardiovasc Res. 2004;62(3):500–8.

88. Lemoine MD, Duverger JE, Naud P, Chartier D, Qi XY, Comtois P, et al. Arrhythmogenic left atrial cellular electrophysiology in a murine genetic long QT syndrome model. Cardiovasc Res. 2011;92(1):67–74.

89. S Ob, Holmes AP, Johnson DM, Kabir SN, C Os, M Or, et al. Increased atrial effectiveness of flecainide conferred by altered biophysical properties of sodium channels. J Mol Cell Cardiol. 2022;166:23–35.

90. Winters J, von Braunmuhl ME, Zeemering S, Gilbers M, Brink TT, Scaf B, et al. JavaCyte, a novel open-source tool for automated quantification of key hallmarks of cardiac structural remodeling. Scientific reports. 2020;10(1):20074.

91. Kavanagh DM, Smyth AM, Martin KJ, Dun A, Brown ER, Gordon S, et al. A molecular toggle after exocytosis sequesters the presynaptic syntaxin1a molecules involved in prior vesicle fusion. Nature communications. 2014;5:14.

92. Ovesny M, Krizek P, Borkovec J, Svindrych ZK, and Hagen GM. ThunderSTORM: a comprehensive ImageJ plug-in for PALM and STORM data analysis and super-resolution imaging. Bioinformatics. 2014;30(16):2389–90.

93. Pike JA, Khan AO, Pallini C, Thomas SG, Mund M, Ries J, et al. Topological data analysis quantifies biological nano-structure from single molecule localization microscopy. Bioinformatics (Oxford, England). 2020;36(5):1614–21.

94. Love MI, Huber W, and Anders S. Moderated estimation of fold change and dispersion for RNA-seq data with DESeq2. Genome Biol. 2014;15(12):550.

95. Jaoude SA, Leclercq JF, and Coumel P. Progressive ECG changes in arrhythmogenic right ventricular disease. Evidence for an evolving disease. European heart journal. 1996;17(11):1717–22.

96. Brembilla-Perrot B, Jacquemin L, Houplon P, Houriez P, Beurrier D, Berder V, et al. Increased atrial vulnerability in arrhythmogenic right ventricular disease. Am Heart J. 1998;135(5 Pt 1):748–54.

97. Peters S, Trummel M, and Meyners W. Prevalence of right ventricular dysplasia-cardiomyopathy in a non-referral hospital. Int J Cardiol. 2004;97(3):499–501.

98. Saguner AM, Ganahl S, Kraus A, Baldinger SH, Medeiros-Domingo A, Saguner AR, et al. Clinical role of atrial arrhythmias in patients with arrhythmogenic right ventricular dysplasia. Circulation journal: official journal of the Japanese Circulation Society. 2014;78(12):2854–61.

99. Wu L, Guo J, Zheng L, Chen G, Ding L, Qiao Y, et al. Atrial Remodeling and Atrial Tachyarrhythmias in Arrhythmogenic Right Ventricular Cardiomyopathy. The American journal of cardiology. 2016;118(5):750–3.

100. Bourfiss M, Te Riele AS, Mast TP, Cramer MJ, JF Vdh, TA Vanv, et al. Influence of Genotype on Structural Atrial Abnormalities and Atrial Fibrillation or Flutter in Arrhythmogenic Right Ventricular Dysplasia/Cardiomyopathy. J Cardiovasc Electrophysiol. 2016;27(12):1420–8.

101. Mazzanti A, Ng K, Faragli A, Maragna R, Chiodaroli E, Orphanou N, et al. Arrhythmogenic Right Ventricular Cardiomyopathy: Clinical Course and Predictors of Arrhythmic Risk. J Am Coll Cardiol. 2016;68(23):2540–50.

102. Gilljam T, Haugaa KH, Jensen HK, Svensson A, Bundgaard H, Hansen J, et al. Heart transplantation in arrhythmogenic right ventricular cardiomyopathy - Experience from the Nordic ARVC Registry. Int J Cardiol. 2018;250:201–6.

103. Wu L, Bao J, Liang E, Fan S, Zheng L, Du Z, et al. Atrial involvement in arrhythmogenic right ventricular cardiomyopathy patients referred for ventricular arrhythmias ablation. J Cardiovasc Electrophysiol. 2018;29(10):1388–95.

104. Mussigbrodt A, Knopp H, Efimova E, Weber A, Bertagnolli L, Hilbert S, et al. Supraventricular arrhythmias in patients with arrhythmogenic right ventricular dysplasia/cardiomyopathy associate with long-term outcome after catheter ablation of ventricular tachycardias. Europace: European pacing, arrhythmias, and cardiac electrophysiology: journal of the working groups on cardiac pacing, arrhythmias, and cardiac cellular electrophysiology of the European Society of Cardiology. 2018;20(7):1182–7.

105. Cardona-Guarache R, Astrom-Aneq M, Oesterle A, Asirvatham R, Svetlichnaya J, Marcus GM, et al. Atrial arrhythmias in patients with arrhythmogenic right ventricular cardiomyopathy: Prevalence, echocardiographic predictors, and treatment. J Cardiovasc Electrophysiol. 2019;30(10):1801–10.

106. Kikuchi N, Shiga T, Suzuki A, and Hagiwara N. Atrial tachyarrhythmias and heart failure events in patients with arrhythmogenic right ventricular cardiomyopathy. IntJ Cardiol Heart Vasc. 2020;31:100669.

