## Supplement for "Male sex hormone and reduced plakoglobin jointly impair atrial conduction and cardiac sodium currents"

### Full methods

**Ethical statement**

All animal procedures were approved by the UK Home Office (PPL number 30/2967 and PFDAAF77F) and by the institutional review board of University of Birmingham. All animal procedures conformed to the guidelines from Directive 2010/63/EU of the European Parliament on the protection of animals used for scientific purposes. Wildtype (WT) and plakoglobin deficient (Plako$^{+/-}$) male 129/Sv mice (1) were housed in individually ventilated cages, (2-7 mice/ cage), under standard conditions: 12 h light/dark circle, 22 °C and 55% humidity. Food and water were available *ad libitum*. The general health status of all mice (bearing, grooming, behaviour) used in the study was monitored daily and immediately prior all animal procedures.

**Chronic 5α-dihydrotestosterone (DHT) exposure and experimental timeline**

Mice were age and weight matched and assigned to either DHT or placebo/control treatment groups. At 8-10 weeks of age, they were fitted with a subcutaneous osmotic mini-pump (Alzet, Cupertino, CA, US) containing either DHT (62.5 mg/mL in ethanol), or ethanol alone (Control - Ctr). At least 40 minutes before pump implantation, mice were subcutaneously injected with 0.05 mL of Buprenorphine. Pump implant was performed under anaesthesia with isoflurane inhalation in $O_2$. After implant, mice were subcutaneously injected with 0.5 mL of glucose saline. An echocardiogram was performed at 6 weeks exposure. At 4 months of age, following the 6 weeks DHT or control treatment, mouse hearts were extracted by thoracotomy under deep terminal anaesthesia (4-5% isoflurane inhalation in $O_2$), and used for *in vitro* experimental analysis, as summarised in Figure 2.

**Echocardiography**

Echocardiography was performed as previously described (2, 3) with a dedicated small animal system (Vevo 2100; Visual Sonics, Toronto, Ont, Canada, now Fujifilm) under light anaesthesia (0.5-2% isoflurane in $O_2$) in a warm environment to keep body temperature stable. Cardiac dimensions and function were assessed in long and short parasternal axis and apical views. Mean values taken from at least 3 images in 2D and M-Mode. Experiment and analysis were performed in a blinded fashion and the analysis was performed by a second blinded observer.

**Steroid measurements**

Serum DHT was determined by ultra-performance liquid chromatography-tandem mass spectrometry (LC-MS/MS) as described (4). In brief, aliquots of samples, calibrator and controls were combined with the internal standard mixture to monitor recovery. All samples were extracted using Oasis MAX SPE system Plates (Waters, Milford, MA, USA). The chromatographic separation was carried out using an ultra performance liquid chromatography (UPLC) system, which is connected to a Quattro Premier/XE triple Quad mass spectrometer (Waters, Milford, MA). A Waters Acquity UPLC BEH C18 column (1.7 µm, 100 x 2.1) was used at a flow rate of 0.4 mL/min at 50C. Water and acetonitril with 0.01% formic acid were used as mobile phase. Two mass transitions were monitored. The following optimized voltages were used: capillary voltage, 3.5 kV, cone voltage 28-33 V, collision energy 18-25 eV, dwell time 0.01-0.08 s depending on the steroid, source temperature 120 °C, desolvation temperature 450 °C. Argon was used as collision gas. Data were acquired with MassLynx 4.1 software and quantification was performed by TargetLynx software (Waters, Milford, MA). During all analyses the ambient temperature was kept at 21°C by air conditioning. Limit of detection: 0.054 nmol/L; Limit of quantification: 0.1 nmol/L, coefficient of variation inter-assay: 4-8% for 0.75 nmol/L and 7.5 nmol/L and intra-assay between 2-3%.

**Left atrial optical mapping**

Murine hearts were mounted on a vertical Langendorff apparatus (Hugo Sachs, March-Hugstetten, Germany) and the aorta was retrogradely perfused at 36-37°C, pH 7.4 with Krebs-Henseleit (KH) solution plus Di-4-ANEPPS (17.5 µM; Cambridge Bioscience, CA, USA). The left atrium (LA) was removed and pinned out in a recording chamber with the superior surface exposed (6, 7). The LA was continuously superfused with a KH solution containing in mM: NaCl 118; $NaHCO_3$ 24.88; $KH_2PO_4$ 1.18; Glucose 5.55; $MgSO_4$ 0.83; $CaCl_2$ 1.8; KCl 3.52, equilibrated with 95% $O_2$/5% $CO_2$, pH 7.4. Blebbistatin (10 µM; Cayman Chemical, Michigan, USA) was added to the superfusate to prevent contraction artefacts. After 15 minutes equilibration, the LA was paced (2 ms duration pulses, 2x diastolic voltage threshold) at 120-80 ms cycle length (CL) using bipolar platinum electrodes and constant voltage stimulator (Digitimer, Welwyn Garden City, UK). Di-4-ANEPPS was excited at 530 nm by four LEDs (Cairn Research, Kent, UK) and fluorescence was captured at 630±20 nm using a second generation, high spatial resolution CMOS ORCA flash 4.0 camera (Hamamatsu Photonics, Japan). Images were captured at a sampling rate of 0.987 kHz (128 x 2048 pixels) and digitised using WinFluor V3.4.9 (Dr John Dempster, University of Strathclyde, UK).

Raw images were used to manually measure LA unfolded area in FIJI. Activation maps, conduction velocity vectors and optical action potentials were generated using MATLAB algorithms as described previously (6-9). Individual beat activation times were calculated from activation maps constructed by measuring the depolarisation midpoint (time of 50% upstroke amplitude) of the optical action potential at each pixel. Percentage tissue activation was then measured as a function of time, with time to 95% activation compared between the individual beats.

**Left atrial transmembrane action potential analysis**

Transmembrane action potentials (TAPs) were recorded from isolated, superfused LA using floating glass microelectrodes (resistance 15-30 MΩ) filled with 3 M KCl (6-8). Preparations were paced incremental from 1000 ms to 80 ms. Voltage was digitised at 20 kHz and was unfiltered. TAPs used for analysis were following 200 stimulations to allow for sufficient rate adaptation.

**RNA isolation, sequencing and analysis**

Left and right atria tissue samples were collected from excised hearts after quick perfusion to rinse out remaining blood. Total RNA was extracted using Direct-zol RNA MiniPrep kit (Zymo Research, Irvine, U.S.A.). RIN values (>7) were checked using the Bioanalyzer RNA 6000 Nano Kit (Agilent, Santa Clara, U.S.A.). mRNA enrichment and subsequent cDNA NGS library preparation was completed using a NEBNext Poly(A) mRNA Magnetic Isolation Module and Ultra II Directional RNA Library Prep Kit for Illumina (New England BioLabs, Hitchin, U.K). NGS library size distribution was determined by use of the Bioanalyzer High Sensitivity DNA Kit (Agilent, Santa Clara, U.S.A.) and quantified using the KAPA Library Quantification Kit (Roche, Basel, Switzerland). Equimolar pooled libraries were sequenced 75 cycles in a single read mode on the NextSeq 500 System (v2.5 Chemistry, Illumina). RNA-seq FASTQ files were aligned on HISAT2 (version 2.1.0) using Ensembl Mus Musculus reference GRCm38 (10, 11). Aligned reads were counted using HTseq version 0.11.2 (12). Required transformations through different RNA-seq analysis steps were done using Samtools version 1.4 (13). Differential expression was obtained using DESeq2 in R (14) (http://www.R-project.org/, R 3.4.1). Ensembl IDs were transformed to gene symbols using BioTools (https://www.biotools.fr/mouse/ensembl_symbol_converter, accessed: 28/02/2020).

Data will be made publically available at https://www.ncbi.nlm.nih.gov/geo/.

**Protein quantification**

Snap-frozen left ventricles were homogenized and resuspended in lysis buffer. 50 µg whole cell lysate per sample were used for SDS-PAGE and semi-dry blotting. Membranes were blocked in 10% milk in TBS-Tween and antibodies (GAPDH, Santa Cruz, sc-365062, 1:300; Connexin 43, Thermo Scientific, 13-8300, 1:250) were diluted in 4% milk in TBS-Tween.

Protein-Antibody-HRP complexes were visualized using ECL substrates on a ChemiDoc Imaging System (Bio-Rad, Hercules, California, USA).

**Left atrial cardiomyocyte cell isolation**

Hearts were mounted onto a Langendorff apparatus and perfused with the following three solutions, equilibrated with 100% $O_2$: (i) HEPES-buffered modified Tyrode's solution containing in mM: NaCl 145, KCl 5.4, $CaCl_2$ 1.8, $MgSO_4$ 0.83, $Na_2HPO_4$ 0.33, HEPES 5, and glucose 11 (pH 7.4, NaOH) x 5 min; (ii) $Ca^{2+}$-free Tyrode's solution × 5 min; (iii) Tyrode's enzyme solution containing 20 µg/mL Liberase™ (Roche, Indianapolis, IN) or a collagenase/protease mix (640 µg/mL collagenase type II, 600 µg/mL collagenase type IV and 50 µg/mL protease (Worthington, Lakewood, NJ)), 20 mM taurine and 30 µM $CaCl_2$ × 15-20 min. The LA was removed and placed into a high-$K^+$ modified Kraft-Bruhe (KB) solution containing in mM: DL-potassium aspartate 10, L-potassium glutamate 100, KCl 25, $KH_2PO_4$ 10, $MgSO_4$ 2, taurine 20, creatine 5, EGTA 0.5, HEPES 5, 0.1% bovine serum albumin (BSA, Sigma), and glucose 20 (pH 7.2, KOH). The LA was dissected into small strips and then cells were released by gentle trituration with fire-polished glass pipettes (1-2 mm diameter). Cells to be used for patch clamp experiments were gradually reintroduced to $Ca^{2+}$ over a period of 2 hours to reach a final concentration of 1.8 mM. All experiments were performed within 4-8 hours of isolation.

**Sodium current recordings**

Isolated atrial cells were plated on laminin coated coverslips and superfused at 3-4 mL/min at 22±0.5 °C with a low sodium external solution containing in mM: NaCl 10, $C_5H_{14}ClNO$ 130, HEPES 10, $CaCl_2$ 1.8, MgCl 1.2, $NiCl_2$ 2, glucose 10, pH 7.4 (CsOH). Peak $I_{Na}$ was recorded in voltage clamp mode using borosilicate glass pipettes (tip resistance 1.5–2.5 MΩ). The internal pipette solution contained in mM: NaCl 5, CsCl 115, HEPES 10, EGTA 10, MgATP 5, $MgCl_2$ 0.5 and TEA 20, pH 7.2 (CsOH). Currents were digitised at 50 kHz and low pass filtered at 20 kHz. Series resistance was compensated between 60 and 80%. Experiments were terminated if series resistance increased abruptly or was greater than 10

MΩ (usually 4-8 MΩ). Current-voltage relationships were examined using 100 ms step depolarisations over test potentials ranging from -95 mV to +40 mV, in 5 mV increments, from a holding potential of -100 mV. I/V curves were fitted using the modified Boltzmann equation: $I_{Na} = G_{max}(V_m – V_{rev})/(1+Exp[(_{V0.5}-V_m)/k])$, where $I_{Na}$ is the current density at an equivalent test potential ($V_m$), $G_{max}$ is the peak conductance (nS), $V_{rev}$ is the reverse potential, $V_{0.5}$ is the membrane potential at 50% current activation and k is the slope constant (15) (16). Time dependent peak $I_{Na}$ recovery kinetics were evaluated using standard 20 ms P1-P2 pulse protocols (-100 mV to -30 mV) with increasing time delay ranging from 1 to 100 ms. Measurements of steady state inactivation of $I_{Na}$, were made by applying 500 ms pre-pulses ranging from -120 mV to -40 mV in 5 mV increments prior to the test potential (-30 mV for 100 ms). Individual atrial cell capacitance was calculated by integrating the cellular capacitance current elicited by 10 mV depolarisation from a holding potential of -80 mV.

**Atrial fiber size and extracellular matrix composition analysis**

Hearts were isolated and coronary arteries cleared with blood using KH buffer, before being snap-frozen in optimal cutting compound (OCT). For fiber size analysis, 10 μm atrial sections were stained with FITC-conjugated wheat germ agglutinin WGA (1:1000) and imaged on Leica DM6000 B fully automated fluorescence microscope (Leica Microsystems, Wetzlar, Germany) with a 40 x objective. Scaled images were extracted using LASX (Leica Application Suite X, Leica Microsystems, Wetzlar, Geramany) imaging suite and cell diameter and endomysial fibrosis were quantified from sections using the JavaCyte ImageJ plugin (17).

**Super-resolution microscopy and $Na_v1.5$ cluster analysis**

LA cells were plated on 10 mm diameter laminin coated coverslips (Mattek, 35 mm dish, 1.5# coverglass) and then fixed in 4% paraformaldehyde for 90 minutes to ensure maximal immobilization of cellular proteins. Cells were permeabilised with a 0.1% Triton in PBS solution for 5 minutes and blocked in a 5% BSA in PBS solution for 60 minutes. The primary polyclonal rabbit anti-Nav1.5 antibody (ASC-005, Alomone Laboratories, Jerusalem, Israel) (1:50) was diluted in blocking solution and incubated overnight at 4°C. Cells were washed three times in PBS, blocked again for 30 minutes and then incubated in secondary antibody fragment (F(ab')2- Goat anti-Rabbit IgG, Alexa Fluor 647, A212-56, 1:1000, ThermoFisher Scientific, Waltham, MA, USA) for 90 minutes at room temperature. Direct stochastic optical reconstruction microscopy (dSTORM) experiments were performed on a NIKON Eclipse Ti inverted N-STORM microscope equipped with a NIKON APO 100 x 1.49 NA total internal reflection fluorescence (TIRF) oil immersion objective. Laser illumination was provided using 405 nm (20 mW), 491 nm (100 mW), 561 mW (100 mW) and 640 nm (200 mW) solid-state lasers. A NIKON Perfect Focus System (PFS) ensured minimal lateral drift during acquisition. Immunolabelled samples were imaged in 0.5 mg/mL glucose oxidase, 40 µg/mL catalase, 10% wt/vol glucose and 100 mM MEA in PBS, pH 7.4 to induce Alexa 647 blinking. During dSTORM acquisition, the sample was continuously illuminated at 640 nm for 20,000 frames, fluorescence emission was filtered through a single band far red emission filter (MBE47200 N-STORM cube) and detected using an Andor iXon Ultra DU897 EMCCD camera (256 x 256 pixels, 160 nm effective pixel size, 9.2 ms exposure time). the angle of illumination as well as the z-depth were kept as consistent as possible across different samples. Samples were maintained in an OKO environmental chamber at 27 $^{o}$C for maximum system stability during imaging. Immunolabelled samples were imaged in 0.5 mg/mL glucose oxidase, 40 µg/mL catalase, 10% wt/vol glucose and 100 mM MEA in PBS, pH 7.4 to induce Alexa 647 blinking.

Fluorescence detections were localised in ThunderSTORM (18) using default settings for the localisation step. Post-localisation the samples were subjected to drift correction (cross correlation), filtering for localizations with a spatial uncertainty below 40 nm, duplicate removal (distance threshold 75 nm) and merging (performed with a maximum distance of 75 nm, maximum off frames 1, maximum frames per molecule 0).

Depending on the overall number of blinking events in a sample, either 20,000 or 5,000 of the post-processed images were subjected to cluster analysis. Due to variability in blinking events across different imaging sessions, resulting cluster parameters from each group were normalised to the mean of the wildtype control group, acquired during the same imaging session.

For cluster analysis, we used a topological analysis tool (5), with codes customized to cardiomyocyte images in an R environment and the following set parameters: r=40 nm, Threshold=10, minimal number of detections per cluster = 10. Prior to cluster analysis, only data points within the cell area were selected using the normalized Ripley's K-function (H-function, $H(r) = L(r)-r$) with a set linking distance of r = 1 µm and only points with $H(r) > 0$, indicating a non-dispersed spatial distribution, were included (background will be randomly distributed and hence $H(r) < 0$). The minimum area to be detected as 'cell' was set to 15 $\mu m^2$. Final super-resolution rendered images were achieved by applying a normalised Gaussian with magnification 50 and the lateral uncertainty set to the calculated uncertainty of the localisation for each blink (20 nm). Cell area selection and cluster map images for visualization were generated using MATLAB.

**Statistics**

Data was first subjected to outlier analysis, ROUT method, based around a false discovery rate, where $\alpha = 0.01$ and outliers were removed (Prism v8, GraphPad Software, La Jolla, CA, USA). Significance was taken at $p<0.05$, applying two-way or two-way repeated measures ANOVA with post hoc t-tests or Bonferroni correction, as appropriate (Prism v8, GraphPad Software, La Jolla, CA, USA). All experiments and analyses were performed blinded to genotype and treatment.

### Supplementary Figures and Tables

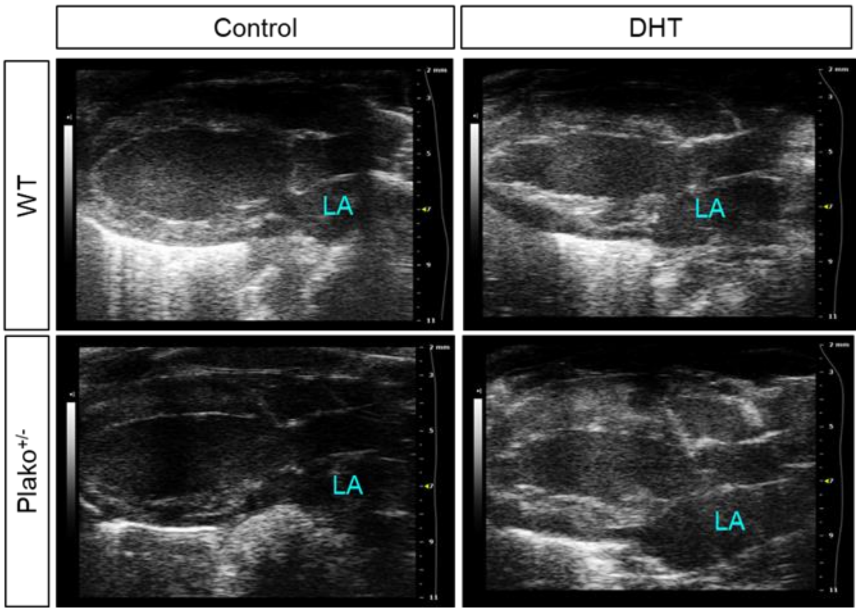


**Supplementary Figure 1 - Echocardiographic assessment of murine hearts**

Example echocardiographic images obtained in parasternal long axis view, displayed during systole (atrial diastole).

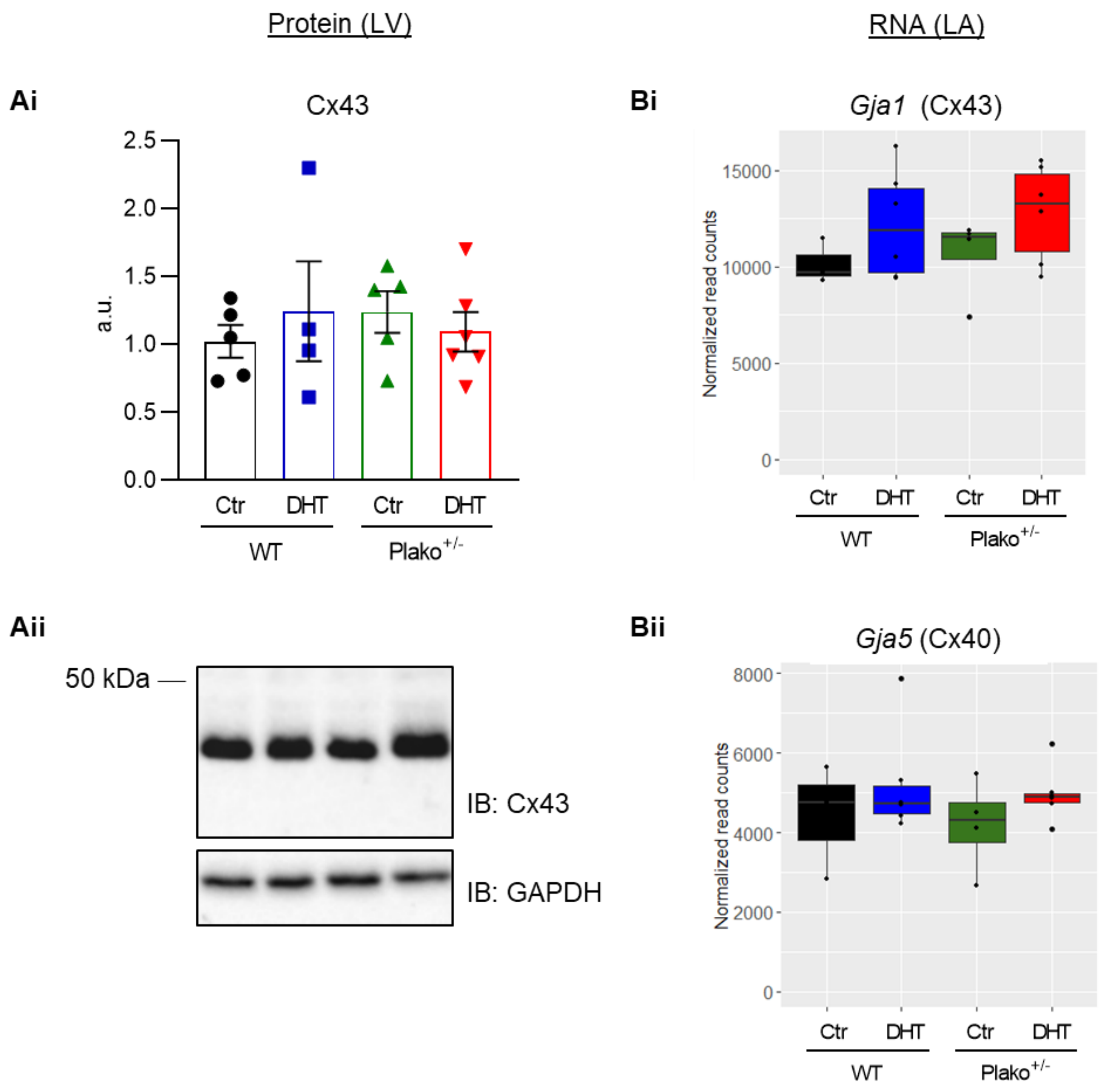


**Supplementary Figure 2 - Expression levels of Connexins**

(**Ai**) Quantification of Cx43 protein expression in LV normalized to the housekeeping protein GAPDH and given as a fraction of the mean expression value in WT Ctr tissue. (**Aii**) Representative Western blot of Connexin 43 (Cx43) in murine left ventricular (LV) tissue.
(**Bi**) Left atrial (LA) RNA expression levels of Gja1, encoding for Connexin 43, Cx43 and of (**Bii**) Gja5, encoding for Connexin 40, Cx40.

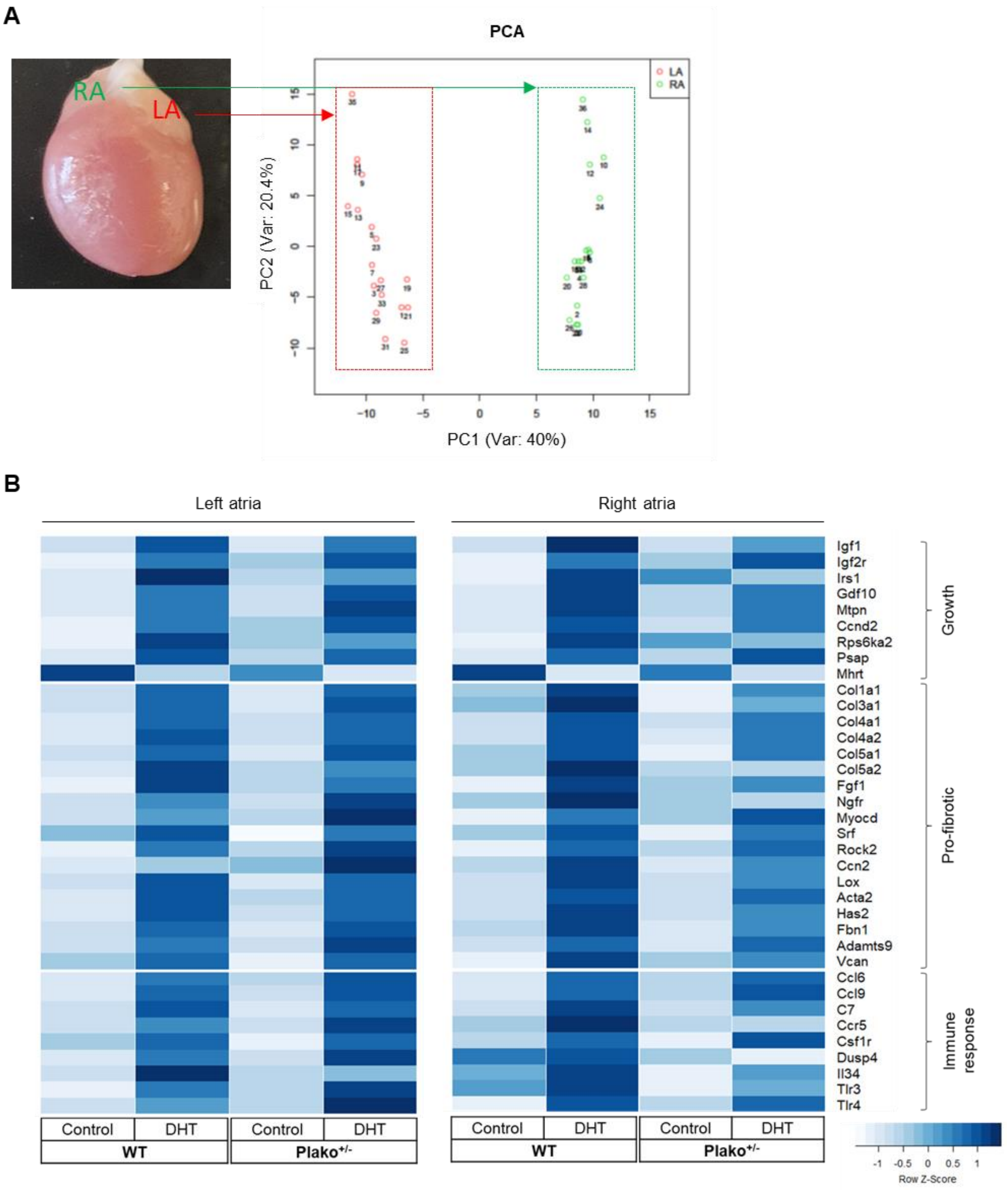


**Supplementary Figure 3 - Atrial RNA expression**

(A) Principal component analysis (PCA) plot of all atrial samples (LA, left atria, red; RA, right atria, green) used for RNA sequencing analysis. The PCA was generated using the top 250 transcripts obtained from the "variance stabilizing transformation" of DESeq2. Range of unique mapped reads/sample across all samples: 16592072 to 22124528. Read length = 75 bp (B) Heat map illustrating differential expression of selected genes in LA and RA resulting from RNA sequencing analysis. n= WT Control: 3 LA / 3 RA; WT DHT: 6 LA / 6 RA; Plako$^{+/-}$ Control: 4 LA / 4 RA; Plako$^{+/-}$ DHT: 6 LA / 6 RA

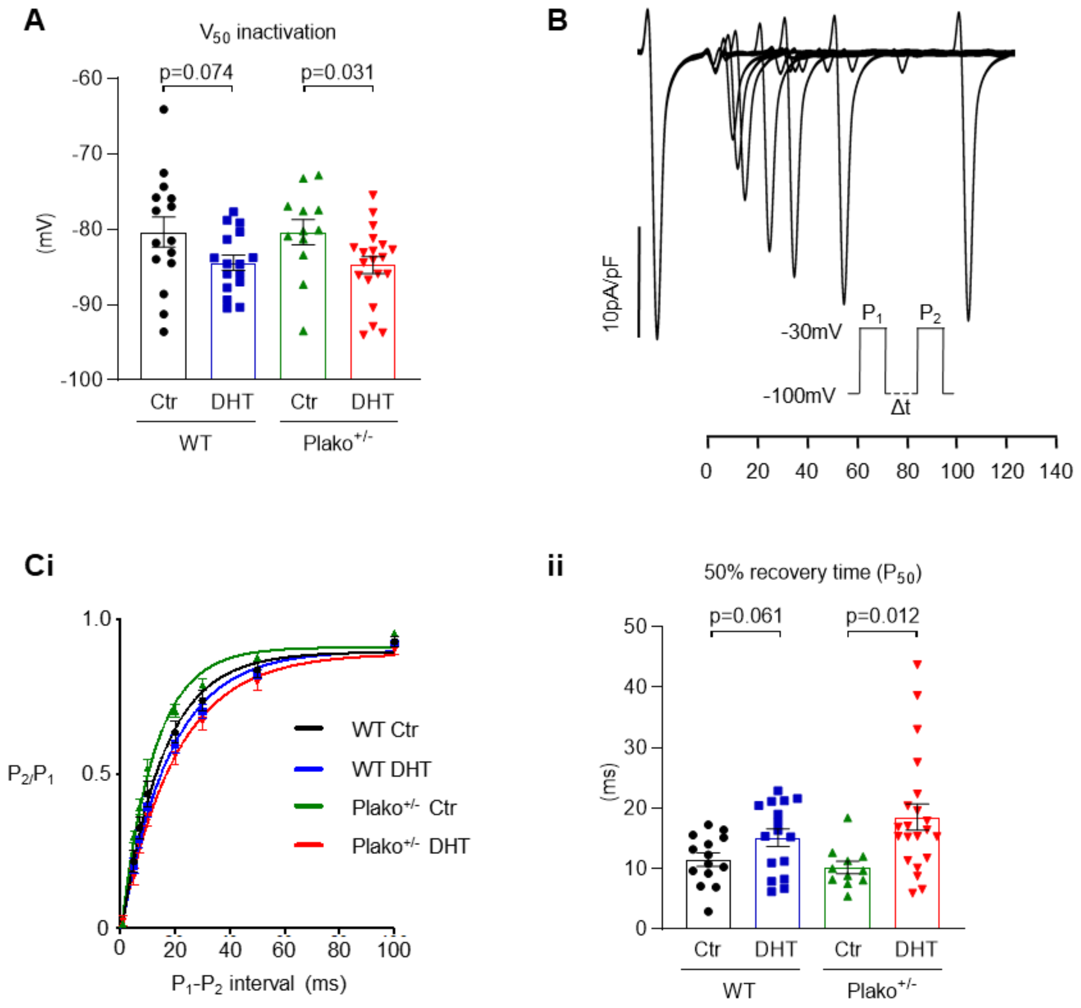


**Supplementary Figure 4 - Sodium channel kinetics**

Exposure to DHT has a significant effect on (**A**) Mean V50 values for steady state inactivation of sodium channels (2-way ANOVA, $p<0.05$). Results of post hoc t-test are indicated on the graph. All data points are shown as well as mean ± SEM (**B**) Example trace showing time dependent sodium channel recovery. (**Ci**) Mean curves showing recovery times. Data plotted as mean ± SEM. DHT treatment has a significant effect on (**Cii**) 50% sodium channel recovery times ($P_{50}$) for each group (two-way ANOVA, $p<0.05$). Results of post hoc t-test are indicated on the graph. Plako$^{+/-}$ LA DHT-treated cells have delayed recovery. All data points are shown as well as mean ± SEM. For C: WT Ctr (n=14 cells, N=5 LA), WT DHT (n=16 cells, N=5 LA), Plako$^{+/-}$ Ctr (n=11 cells, N=4 LA) and Plako$^{+/-}$ DHT (n=21 cells, N=5 LA).

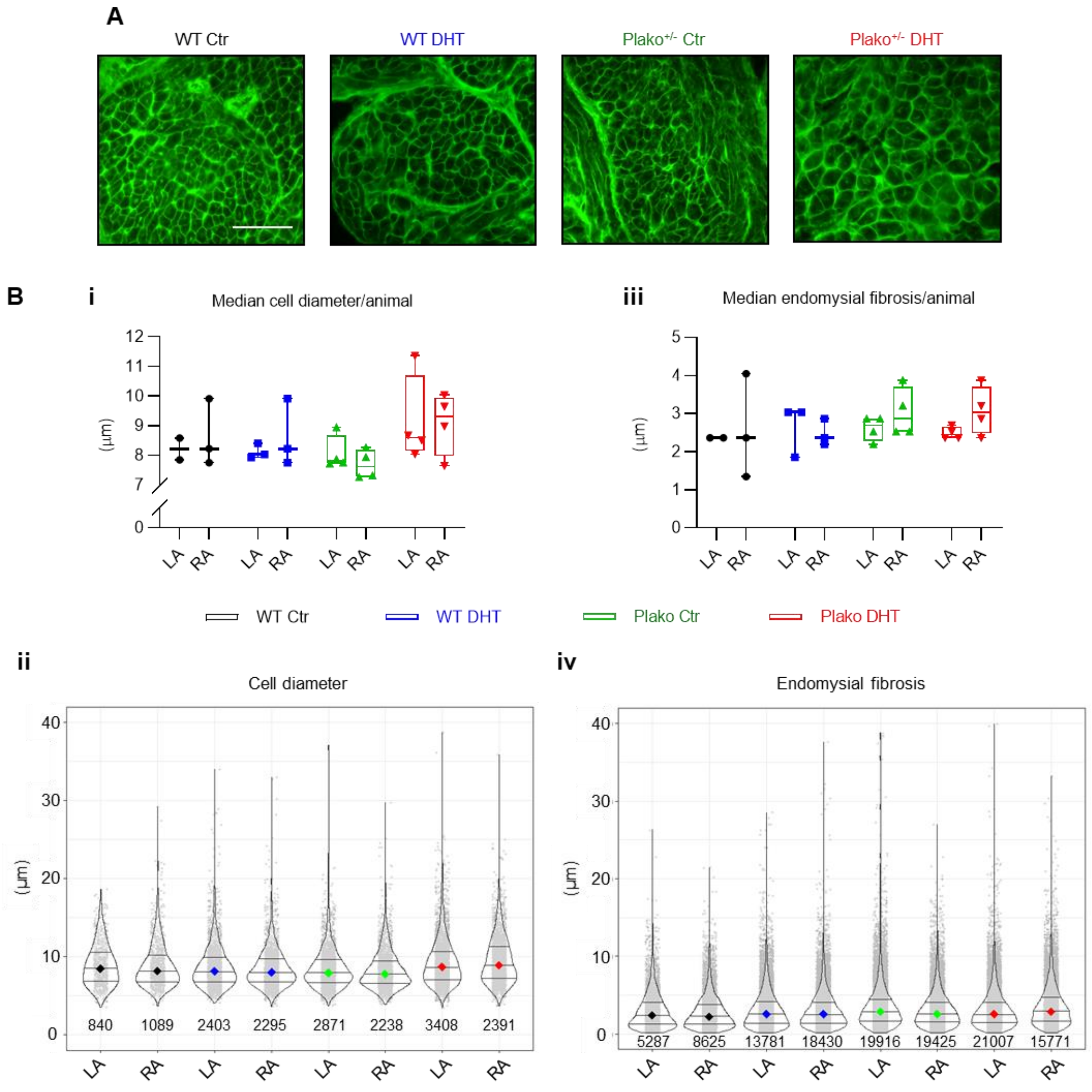


**Supplementary Figure 5 - Histological analysis of atria**

(**A**) Example left atrial (LA) sections stained with FITC-conjugated wheat germ agglutinin (lectin, 400 x magnification) for automated quantification of (**Bi&ii**) cell diameter and (**Biii&iv**) endomysial fibrosis (cell-cell-distance) in both, LA and right atria (RA). Scale bar 20 µm
(Bi and Biii) Single data points denote medians of individual measurements per atrium
(Bii and Biv) individual measurements data points (light grey) with violin plots including medians (color-coded per group) and IQR. (Bii) IQR (µm) are as follows: WT Placebo LA: 3.678, WT Placebo RA: 3.248, WT DHT LA: 3.1375, WT DHT RA: 2.89, Plako$^{+/-}$ Placebo LA: 2.8395, Plako$^{+/-}$ Placebo RA: 2.8545, Plako$^{+/-}$ DHT LA: 3.5878, Plako$^{+/-}$ DHT RA: 4.071 (Biv) WT Placebo LA: 2.867, WT Placebo RA: 2.53, WT DHT LA: 2.698, WT DHT RA: 2.698, Plako$^{+/-}$ Placebo LA: 2.698, Plako$^{+/-}$ Placebo RA: 2.529, Plako$^{+/-}$ DHT LA: 2.529, Plako$^{+/-}$ DHT RA: 3.036. Number of cell diameters / distances measured for each group is reported in the respective column of the graphs. Linear mixed modelling revelaed no significant differences between genotype or treatment groups.

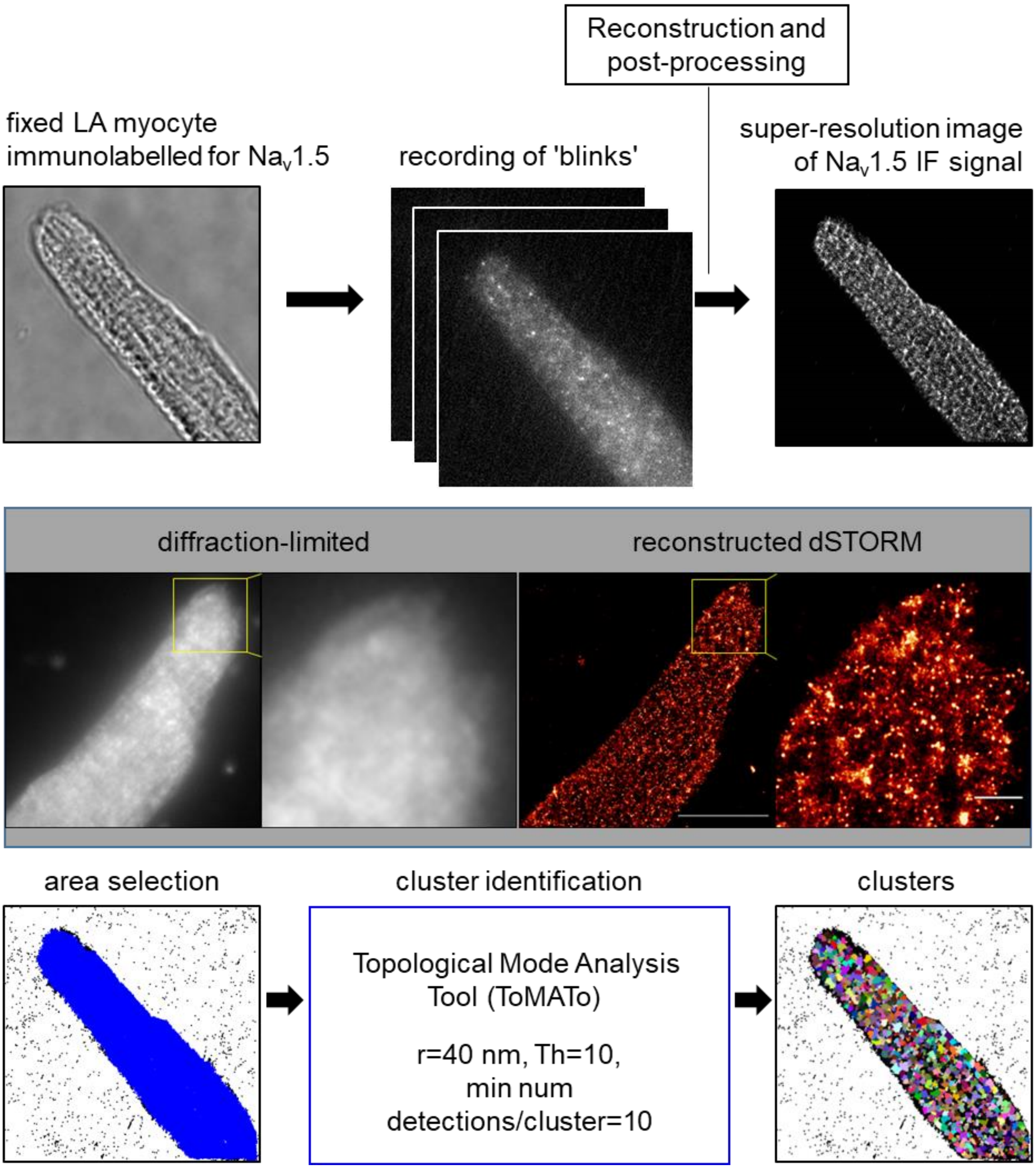


**Supplementary Figure 6 - Workflow for direct stochastic optical reconstruction microscopy (dSTORM) and subsequent clustering analysis in left atrial (LA) cardiomyocytes**

Fixed LA cardiomyocytes were immunolabelled for Na$_V$1.5 and blinking events (blinks) in STORM buffer were recorded. Recordings over time were then post-processed and super-resolution images were reconstructed and false-coloured. Increase in resolution over the diffraction-limited image is depicted in the middle panel. The cell area was selected (blue) and contained detections were subjected to clustering analysis, using a topological approach (5). Proposed clusters are depicted in arbitrary colours, detections not allocated to any cluster remain black.

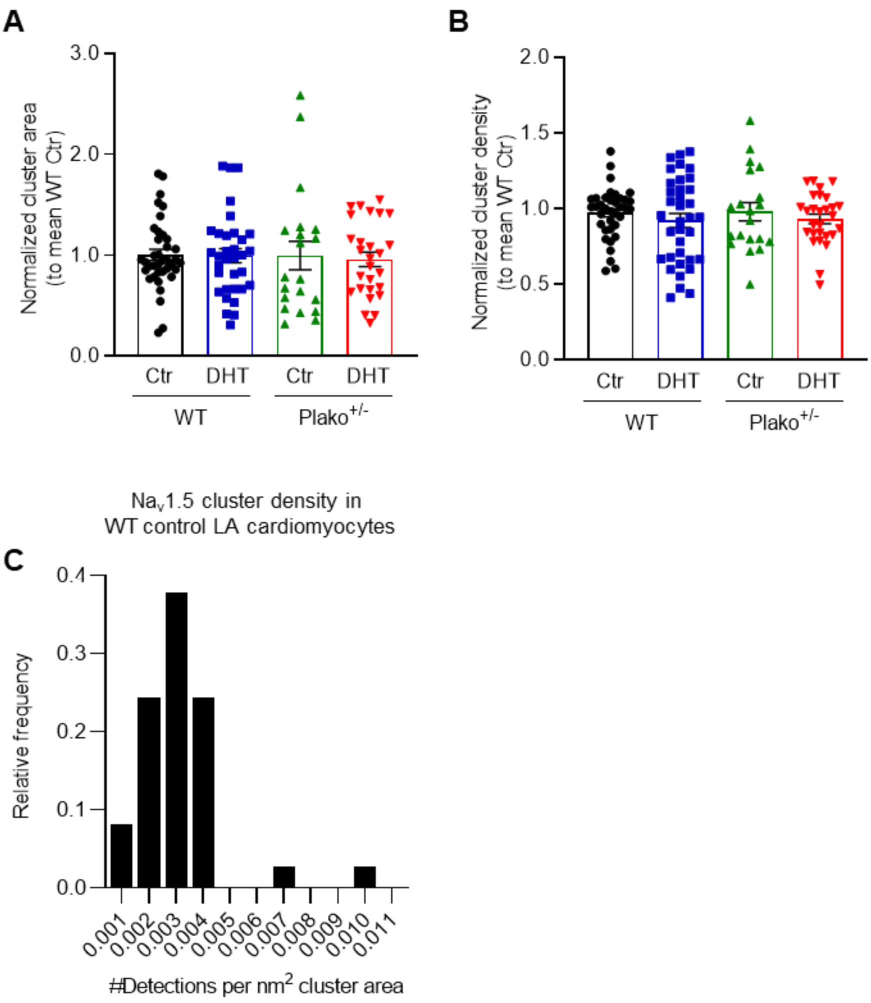


**Supplementary Figure 7 - Sodium channel cluster characteristics**

Neither plakoglobin heterozygous knockout nor exposure to DHT have an effect on (**A**) normalized $Na_v1.5$ cluster area and (**B**) cluster density. (**C**) Relative frequency of $Na_v1.5$ cluster densities (number of detections per $nm^2$ cluster area in WT control left atrial cardiomyocytes (n=38 cells, N=7 LA)).

#### Supplementary Table 1 - Transmembrane action potential analysis

| | WT Control (n=21, N=7) | | | WT DHT (n=13, N=5) | | | Plako$^{+/-}$ Control (n=20, N=7) | | | Plako$^{+/-}$ DHT (n=23, N=8) | | |
|---|---|---|---|---|---|---|---|---|---|---|---|---|
| **Pacing CL (ms)** | **120** | **100** | **80** | **120** | **100** | **80** | **120** | **100** | **80** | **120** | **100** | **80** |
| **RMP (mV)** | -72 ±1 | -71 ±1 | -70 ±1 | -71 ±1 | -70 ±1 | -69 ±1 | -71 ±1 | -70 ±1 | -69 ±1 | -69 ±1* | -68 ±1* | -67 ±1* |
| **APA (mV)** | 84 ±1 | 82 ±1 | 79 ±1 | 82 ±2 | 79 ±2 | 75 ±2 | 82 ±2 | 80 ±2 | 76 ±2 | 74 ±2*#+ | 71 ±2*#+ | 67 ±2*#+ |
| **dV dt$^{-1}$ peak (V/s)** | 118 ±5 | 112 ±5 | 104 ±5 | 120 ±7 | 113 ±7 | 103 ±6 | 116 ±4 | 109 ±4 | 98 ±4 | 89 ±5*#+ | 83 ±5*#+ | 74 ±5*#+ |
| **APD50 (ms)** | 7.1 ±0.4 | 6.8 ±0.4 | 6.4 ±0.4 | 6.5 ±0.3 | 6.1 ±0.3 | 5.7 ±0.4 | 7.2 ±0.4 | 6.8 ±0.4 | 6.5 ±0.4 | 6.8 ±0.2 | 6.6 ±0.2 | 6.3 ±0.2 |
| **APD70 (ms)** | 10.8 ±0.6 | 10.2 ±0.5 | 9.6 ±0.5 | 9.9 ±0.6 | 9.2 ±0.5 | 8.5 ±0.4 | 11.2 ±0.8 | 10.6 ±0.7 | 9.9 ±0.6 | 10.5 ±0.4 | 10.0 ±0.4 | 9.5 ±0.4 |
| **APD90 (ms)** | 20.5 ±1.0 | 19.2 ±0.8 | 17.9 ±0.8 | 19.2 ±1.0 | 17.7 ±0.9 | 15.9 ±0.8 | 21.4 ±1.4 | 20.0 ±1.3 | 18.5 ±1.1 | 20.2 ±0.7 | 19.1 ±0.6 | 17.6 ±0.6 |
| **AT (ms)** | 6 ±0.3 | 7 ±0.4 | 7 ±0.4 | 7 ±0.5 | 7 ±0.5 | 8 ±0.6 | 7 ±0.3 | 8 ±0.4 | 8 ±0.4 | 9 ±0.8 *#+ | 10 ±1.0 *#+ | 11 ±1.4 *#+ |

Action potentials from microelectrode recordings in paced, superfused left atria. WT, wildtype; DHT, 5α-dihydrotestosterone; CL, cycle length; RMP, resting membrane potential; APA, action potential amplitude; APD50-90, action potential duration at 50-90% repolarisation; AT, activation time. Data presented as mean ±SEM. *$p<0.05$ vs WT Control, $^{\#}p<0.05$ vs WT DHT, $^{+}p<0.05$ vs Plako$^{+/-}$ Control; two-way ANOVA with Bonferroni post-hoc analysis. n=number of measurements, N=number of left atria

**Supplementary Table 2 - Optical action potential analysis**

| | **WT Control (N=6)** | **WT DHT (N=7)** | **Plako$^{+/-}$ Control (N=8)** | **Plako$^{+/-}$ DHT (N=9)** |
|---|---|---|---|---|
| **Pacing CL (ms)** | **80** | **80** | **80** | **80** |
| **APD90 (ms)** | 20.7 ±0.9 | 22.5 ±1.4 | 21.7 ±1.1 | 22.8 ±1.4 |
| **B-to-b Δ APD90 (ms)** | 1.6 ±0.3 | 3.2 ±0.7 | 1.9 ±0.2 | 1.3 ±0.4 |

Action potentials recorded optically from paced, superfused left atria (LA). WT, wildtype; DHT, 5α-dihydrotestosterone; CL, cycle length; APD90, action potential duration at 90% repolarisation; B-to-b Δ APD90, Beat-to-beat difference in APD90. Data presented as mean ±SEM. N=number of LA

#### Supplementary References


1. Kirchhof P, et al. Age- and training-dependent development of arrhythmogenic right ventricular cardiomyopathy in heterozygous plakoglobin-deficient mice. *Circulation.* 2006;114(17):1799-806.
2. Fabritz L, et al. Load-reducing therapy prevents development of arrhythmogenic right ventricular cardiomyopathy in plakoglobin-deficient mice. *Journal of the American College of Cardiology.* 2011;57(6):740-50.
3. Kirchhof P, et al. PITX2c is expressed in the adult left atrium, and reducing Pitx2c expression promotes atrial fibrillation inducibility and complex changes in gene expression. *Circulation Cardiovascular genetics.* 2011;4(2):123-33.
4. Kulle AE, et al. A novel ultrapressure liquid chromatography tandem mass spectrometry method for the simultaneous determination of androstenedione, testosterone, and dihydrotestosterone in pediatric blood samples: age- and sex-specific reference data. *J Clin Endocrinol Metab.* 2010;95(5):2399-409.
5. Pike JA, et al. Topological data analysis quantifies biological nano-structure from single molecule localization microscopy. *Bioinformatics (Oxford, England).* 2020;36(5):1614-21.
6. Yu TY, et al. An automated system using spatial oversampling for optical mapping in murine atria. Development and validation with monophasic and transmembrane action potentials. *Progress in biophysics and molecular biology.* 2014.
7. Holmes AP, et al. A Regional Reduction in Ito and IKACh in the Murine Posterior Left Atrial Myocardium Is Associated with Action Potential Prolongation and Increased Ectopic Activity. *PloS one.* 2016;11(5):e0154077.
8. Syeda F, et al. PITX2 Modulates Atrial Membrane Potential and the Antiarrhythmic Effects of Sodium-Channel Blockers. *Journal of the American College of Cardiology.* 2016;68(17):1881-94.
9. O'Shea C, et al. ElectroMap: High-throughput open-source software for analysis and mapping of cardiac electrophysiology. *Scientific reports.* 2019;9(1):1389.
10. Kim D, et al. Graph-based genome alignment and genotyping with HISAT2 and HISAT-genotype. *Nat Biotechnol.* 2019;37(8):907-15.
11. Zerbino DR, et al. Ensembl 2018. *Nucleic Acids Res.* 2018;46(D1):D754-D61.
12. Anders S, et al. HTSeq--a Python framework to work with high-throughput sequencing data. *Bioinformatics (Oxford, England).* 2015;31(2):166-9.
13. Li H, et al. The Sequence Alignment/Map format and SAMtools. *Bioinformatics (Oxford, England).* 2009;25(16):2078-9.
14. Love MI, et al. Moderated estimation of fold change and dispersion for RNA-seq data with DESeq2. *Genome Biol.* 2014;15(12):550.
15. Spencer CI, et al. Actions of pyrethroid insecticides on sodium currents, action potentials, and contractile rhythm in isolated mammalian ventricular myocytes and perfused hearts. *J Pharmacol Exp Ther.* 2001;298(3):1067-82.
16. Ackers-Johnson M, et al. A Simplified, Langendorff-Free Method for Concomitant Isolation of Viable Cardiac Myocytes and Non-Myocytes from the Adult Mouse Heart. *Circulation research.* 2016.
17. Winters J, et al. JavaCyte, a novel open-source tool for automated quantification of key hallmarks of cardiac structural remodeling. *Scientific reports.* 2020;10(1):20074.

18. Ovesny M, et al. ThunderSTORM: a comprehensive ImageJ plug-in for PALM and STORM data analysis and super-resolution imaging. *Bioinformatics (Oxford, England).* 2014;30(16):2389-90.
